# Microvascular Specificity of Spin Echo BOLD fMRI: Impact of EPI Echo Train Length

**DOI:** 10.1101/2023.09.15.557938

**Authors:** T.W.P. van Horen, J.C.W. Siero, A.A. Bhogal, N. Petridou, M.G. Báez-Yáñez

## Abstract

A spatially specific fMRI acquisition requires specificity to the microvasculature that serves active neuronal sites. Macrovascular contributions will reduce the microvascular specificity but can be reduced by using spin echo (SE) sequences that use a π pulse to refocus static field inhomogeneities near large veins. The microvascular specificity of a SE-echo planar imaging (SE-EPI) scan depends on the echo train length (ETL)-duration, but the dependence is not well-characterized in humans at 7T. To determine how microvascular-specific SE-EPI BOLD is in humans at 7T, we developed a Monte Carlo voxel model that computes the signal of a proton ensemble residing in a vasculature subjected to a SE-EPI pulse sequence. We characterized the ETL-duration dependence of the microvascular specificity by simulating the BOLD signal as a function of ETL, the range adhering to experimentally realistic readouts. We performed a validation experiment for our simulation observations, in which we acquired a set of SE-EPI BOLD time series with varying ETL during a hyperoxic gas challenge. Both our simulations and measurements show an increase in macrovascular contamination as a function of ETL, with an increase of 30% according to our simulation and 60% according to our validation experiment between the shortest and longest ETL durations (23.1 - 49.7 ms). We conclude that the microvascular specificity decreases heavily with increasing ETL-durations. We recommend reducing the ETL-duration as much as possible to minimize macrovascular contamination in SE-EPI BOLD experiments. We additionally recommend scanning at high resolutions to minimize partial volume effects with CSF. CSF voxels show a large BOLD response, which can be attributed to both the presence of large veins (high blood volume) and molecular oxygen-induced *T*_1_-shortening (significant in a hyperoxia experiment). The magnified BOLD signal in a GM-CSF partial volume voxel reduces the desired microvascular specificity and, therefore, will hinder the interpretation of functional MRI activation patterns.

## 1. Introduction

The BOLD contrast is the most common functional MRI (fMRI) contrast mechanism and enables the study of brain function through neurovascular coupling.^1^Neurovascular coupling constitutes a complicated interplay of changes in blood volume, blood flow, and oxygen consumption.^2^The BOLD contrast probes change in oxygen consumption by treating deoxyhemoglobin (dHb) as an endogenous contrast agent and tracking its dynamics.^3^dHb is paramagnetic and alters the local magnetic susceptibility of blood. Water protons diffuse through the magnetic field gradients in and around dHb-containing vessels and lose phase coherence on a voxel scale.^4^Hence, dHb causes a signal decrease. Upon brain activation, cerebral blood flow generally overcompensates for increases in oxygen consumption, thus decreasing the local dHb concentration by the inflow of oxygen-rich arterial blood.^5^This dHb decrease shows as a BOLD signal increase.^6,7^

Confounding vascular-physiological and MR-physics processes reduce spatial specificity and hamper direct inference of neuronal activity from the BOLD signal. The BOLD signal and its spatiotemporal characteristics do not directly reflect neuronal activity, as neurovascular coupling comprises much more than solely local changes in dHb concentration.^4^Direct inference becomes especially difficult at high spatial resolutions, where the spatial specificity is potentially worse than the resolution. Acquiring higher-resolution fMRI images, therefore, does not guarantee a corresponding increase in specificity to the underlying functional activity.^8^At 7T, submillimeter resolutions are achievable, and understanding the factors reflected in the BOLD signal becomes critical.^9^

One of these factors is the underlying vasculature, which constrains the spatial specificity of the BOLD signal.^8^A grid-like pial vasculature supplies blood to and drains blood from the microvasculature - which serves active neuronal sites - via penetrating and ascending vessels.^10^Whereas the microvasculature gives the most spatially specific BOLD contributions, draining pial veins are the largest source of dHb and lead to the strongest and furthest reaching BOLD effect, reducing spatial specificity of the BOLD signal.^8,9,11^Therefore, microvascular specificity - high sensitivity to microvessels and low sensitivity to macrovessels (mainly pial veins) - is desired to infer activation patterns from the fMRI data. The supralinear increase of microvascular contributions with increasing field strength versus a linear increase of macrovascular contributions has been a significant drive to build high field (*≥* 7T) MRI systems with a promise of increased microvascular specificity.

Spin echo (SE) sequences can further reduce macrovascular contamination and thus improve the spatial specificity of the BOLD signal.^4,9,10,12^The refocusing π pulse typical to an SE sequence can partly refocus extravascular 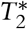 -weighted signals around large veins, where the water protons are approximately static compared to the spatial scale of dHb-induced field inhomogeneities.^3,10^In practice, functional SE sequences often use echo planar imaging (EPI) readouts; EPI readouts implement single-shot 2D acquisitions, which fulfill the high temporal resolution demands required to track activation dynamically for neuroscientific applications.^1^However, the long EPI readout window (‘ETL-duration’ in ms) re-introduces some macrovascular 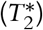 weighting, as full refocusing and *T*^2^-contrast occurs only at the instantaneous spin echo timing.

The dependence of the microvascular specificity on the SE-EPI EsTL-duration is not well-characterized in humans at 7T. Previously, Goense et al. have used laminar fMRI of the macaque brain at 4.7T to establish a positive relationship between readout window size and superficial bias, which is associated with macrovascular contamination.^3^Nevertheless, these experimental results cannot be directly translated to human studies at 7T. Alternatively, biophysical models can be used to study the BOLD behavior at various field strengths and for different vascular properties and pulse sequences. However, previous simulation studies simplified the BOLD signal formation^1^or focused on gradient echo (GE) or SE sequences with instantaneous readouts, i.e. not EPI readouts that are commonly used in fMRI studies.^4,9,10,12,13^

In this work, we bridged this gap by developing a biophysical model that combines the stochastic behavior of water protons with a realistic pulse sequence design for EPI. The model was applied to SE-EPI BOLD acquisitions to give a physiologically-informed recommendation on the sequence design.

Our research question was: “How microvascular specific is SE-EPI BOLD of the human brain at 7T?”

We set out to answer this question for SE-EPI by varying the ETL-duration using simulations and in-vivo measurements. For this purpose, we developed a Monte Carlo voxel model that solves the magnetization behavior of a proton ensemble residing in a vasculature subject to a SE-EPI sequence. We performed a hyperoxic gas challenge experiment at 7T to validate our simulations.

## 2. Methods

### 2.1 Rationale

In this study, we chose a hyperoxic gas challenge to induce a global dHb change as a proxy for functional activation. During a hyperoxic gas challenge, a dedicated gas system (RespirAct™, Thornhill Research, Toronto, CAN) increases the inspired oxygen level above atmospheric concentrations. The following excess of oxygen in the arterial blood plasma reaches the microvasculature and diffuses into the tissue preferentially to hemoglobin-bound oxygen.^14^Thus, similarly to functional activation, administering hyperoxic gas should decrease the local dHb concentration and yield a positive BOLD contrast.^15^Contrary to functional activation, the oxygenation levels should be elevated everywhere in the brain, resulting in a relatively spatially homogeneous BOLD response. We scanned a single healthy subject and included all cortical voxels in our analysis based on the previous assumption.

An additional motivation for using hyperoxia was the lack of expected vasoresponse. Hyperoxia is presumed to induce minimal changes in cerebral blood volume (CBV) given the sensitivity of MRI measurements to this parameter,^10,14^which allowed us to simplify our simulations, disregarding volume changes altogether. During functional experiments, neurovascular coupling induces a CBV increase primarily in large veins and small arteries, which are much more capable at dilation than microvessels.^16^Consequently, our hyperoxia experiment might underestimate macrovascular contamination - and over-estimate the microvascular specificity - compared to an otherwise equivalent functional experiment.

To probe the microvascular specificity, we focused on gray matter (GM) and cerebrospinal fluid (CSF) regions of interest (ROIs), with sulci-located CSF serving as an internal reference devoid of any microvascular contributions. We chose these regions because of their distinct vascular organizations. GM, typically the ROI in fMRI studies, consists of microvessels and larger penetrating and ascending vessels (macrovasculature). CSF does not contain microvessels but encloses large feeding and draining pial vessels (macrovessels). The magnetic field inhomogeneities induced by pial veins often extend into GM voxels. We assume that when the macrovascular contamination decreases, the average CSF BOLD signal will decrease more than its pure GM counterpart. Thus, *we consider the ratio CSF to GM BOLD signal a proxy for macrovascular contamination, with a low CSF/GM BOLD signal ratio indicating a low level of macrovascular contamination and presumably high microvascular specificity*.

We varied the ETL-duration through SENSE (SENSitivity Encoding) acceleration in the phase encoding direction.^17^Increasing the SENSE factor in EPI skips k-space lines, reducing the echo train length (ETL). Provided a constant gradient strength and duration, the ETL-duration in ms is proportional to the ETL: ETL-duration = ETL *×* k-line readout duration. SENSE acceleration reduces the ETL while preserving the image resolution. The resulting k-space undersampling is corrected retrospectively in the SENSE reconstruction but comes at the cost of SNR. Noise breakthrough upon increased undersampling (acceleration) poses a lower limit on the ETL (upper limit on the SENSE factor). The echo time (TE) determines the time interval in which k-lines can be acquired and was constant to ensure a similar image contrast for all ETL values. The upper ETL limit is thus the maximum number of k-space lines that fit within this interval.

### 2.2 Biophysical model

In this section, we provide a general description of our biophysical model. In Section 2.3.1, the simulated GM and CSF physiological parameters (Table C.1) will be specified, along with the pulse sequence (Table C.2) and discretization parameters (Table C.3). In Section 2.3.2, we present additional model details, considerations, and assumptions.

Our biophysical model simulates the mesoscopic tissue-vascular environment within a single, axis-aligned cuboid voxel at the image-space origin. We fill the voxel with vessels represented as infinite cylinders of uniform susceptibility. We add cylinders to the voxel until we reach a target volume fraction *f* .^12^Alternatively, we use a fixed number (*N*) of cylinders.

Water protons walk randomly in the extravascular space. We place each water proton at a random extravascular initial location and add a random displacement vector to the proton location at each discrete time step. We sample this displacement vector 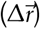 from a normal distribution with zero mean:

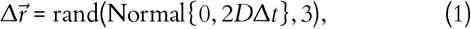

where *D* is the diffusion constant of a proton in the extravascular space and Δ*t* is the discretization time step.^12^The proton undergoes elastic collisions at the vessel and voxel boundaries to prevent passage into the intravascular space or exit from the voxel volume.^9,18^

To efficiently detect proton-boundary interactions, we use a set of bounding volumes, similar to most modern raytracers;^19^instead of checking all interactions between *all* protons and *all* boundaries at *each* time step (billions of computations even for our single voxel simulation), we subdivide the simulation space into a collection of axis-aligned cuboid volumes. We achieve this by splitting up the original voxel volume *n* times in all directions, yielding 8^*n*^ subvolumes. We use these subvolumes to narrow the search space for proton-boundary collisions. At each time point, we know the occupied subvolume and all its contained objects (vessels, voxel walls, and the boundaries of the subvolume itself). When the proton takes its random step, we compute the (potential) sequence of collisions with the contained objects. If a proton hits a subvolume boundary, it freely passes, and we start looking for interactions with objects contained in the new subvolume. And so on, we zoom in on the proton over time and speed up the restricted random walk without sacrificing accuracy. The most time-efficient *n* depends on the complexity of the voxel environment and reflects the trade-off between configuring 8^*n*^subvolumes and reducing the search space.^19^

During its random walk, we integrate the magnetization evolution 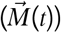 of the proton. At the start of the random walk, the steady-state magnetization is excited by a π/2 pulse. At half the echo time, a π pulse refocuses the magnetization. Throughout the random walk, the magnetization simultaneously relaxes according to the NMR relaxation times (*T*_1_ and *T*_2_) of the environment and accumulates phase due to the magnetic field offset evolution. At each time point *t*, this can be formulated as the matrix multiplication for the Bloch equations:

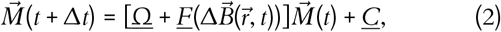

where 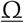 represents RF pulse action and 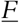 and 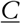 comprise NMR relaxation and precession at a field offset 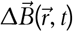. 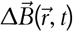 consists of a mesoscopic field contribution and an encoding gradient contribution. Appendix A provides a more detailed description of the Bloch matrices.

We calculate the mesoscopic magnetic field experienced by the proton as a superposition of the magnetic field offsets (caused by non-zero dHb/non-unity oxygenation levels) from sssall the vessels in the voxel evaluated at the proton location:

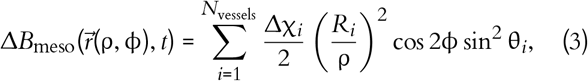

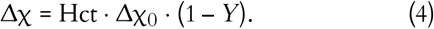

The proton position is expressed in polar coordinates ρ=ρ(*t*) and φ=φ(*t*). Each vessel (cylinder) is characterized by a radius *R*, an azimuthal (θ) and polar (η) angle defining the orientation of the cylinder axis, and a susceptibility shift Δχ (SI units) to the extravascular space. Δχ is linearly proportional to the hematocrit level Hct, the susceptibility difference between fully oxygenated and deoxygenated red blood cells Δχ_0_, and the oxygenation level *Y*.

We compute contributions of the image encoding gradients to the field offset as the inner product of the instantaneous gradient strength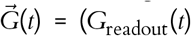, 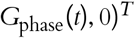 and the proton location 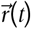 The gradient EPI readout train and phase encoding blips are each simulated by convolving a Dirac comb (setting the gradient strength, polarity, and timings) with a discretized block function with a length equal to the gradient duration for one k-line.

We sample the voxel signal at each EPI gradient echo by computing the ensemble-averaged complex magnetization vector. This average captures the intravoxel dephasing of the transverse magnetization vectors for all protons. The absolute signal is then calculated at each time point and summed to calculate the total signal energy in image space (Parseval’s theorem states that this is equal in k-space and image space). We simulate the proton ensemble in two physiological states: resting (baseline) and active (mimicking the hyperoxic stimulation). We compute the BOLD effect as the percentage absolute signal difference upon a change from resting to active state.

### 2.3 Simulation

#### 2.3.1 Settings

Table C.1 lists all physiological values used in this study. We described the GM vasculature as a combination of microvessels and a single ascending vein. We simulated a single large pial vein in CSF and accounted for an increase in diffusion within CSF compared to GM tissue. We based the relaxation times and oxygenation levels on our hyperoxic stimulus (Figure 3) and included the expected *T*1-shortening effect in CSF upon activation. Figure 1 illustrates the resulting GM and CSF voxel.

**Figure 1.**
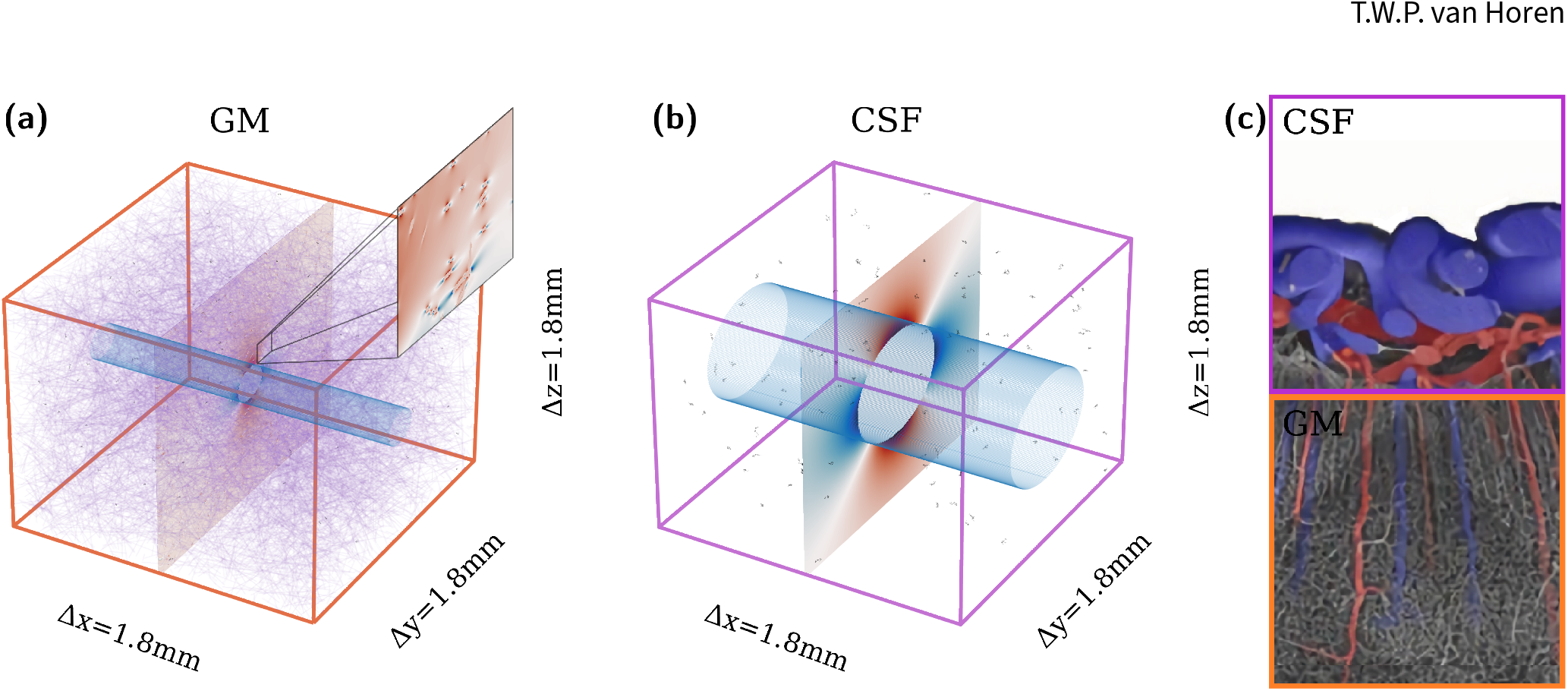
Illustration of the biophysical model. Panel (a) and (b) show the GM and CSF voxel with microvessels indicated in purple and veins indicated in blue. The ascending vein in GM is much smaller than the pial vein in CSF, which complies with the realistic vascular morphology illustrated in panel (c). In this example, the veins are oriented perpendicular to 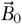 The mesoscopic field offset in the active state is shown for each voxel in a slice perpendicular to the vein. The spatial extent of the field inhomogeneities scales with vessel size, and it is visible on the GM zoom that the microvessels generate small, highly localized field offsets compared to the veins. In black, 100 exemplary proton diffusion paths are plotted, which show the difference in diffusion constant between CSF and GM tissue (3 ×). The relatively high diffusion constant in CSF enhances the sensitivity of the BOLD signal to the pial vein, as refocusing is less effective in more dynamic diffusion regimes. The vascular morphology in panel (c) corresponds to the GM-CSF border within the visual cortex of a macaque (adapted from [20]) and represents a 1.8 × 3.6 mm^2^region.

We matched the MRI pulse sequence parameters to our scan protocol (described in Section 2.4.2); Table C.2 lists all sequence parameters, and Figure 2 shows two exemplary pulse sequences, corresponding to ETL=33 and ETL=71.

**Figure 2.**
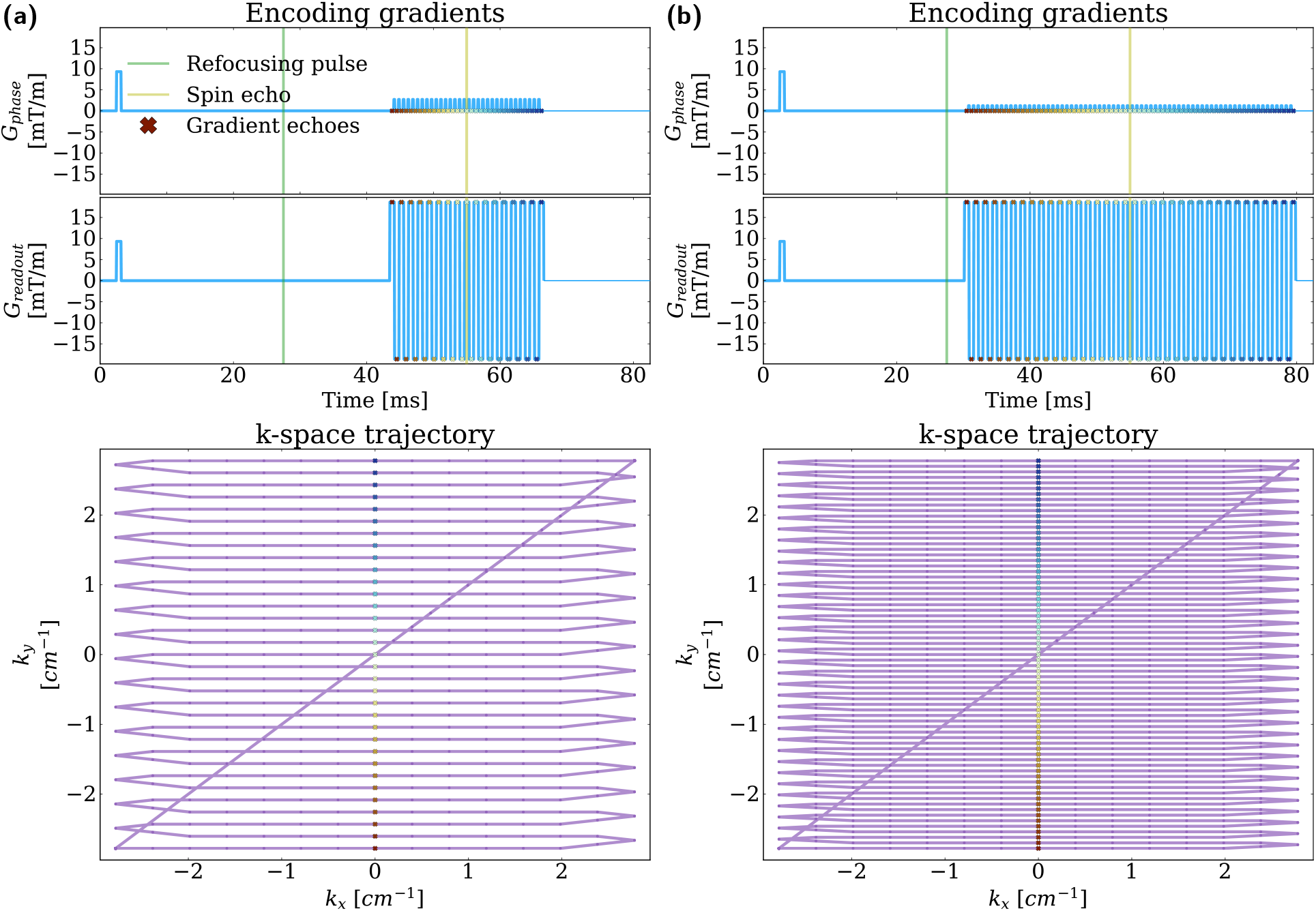
SE-EPI sequence simulated for ETL=33 (a) and ETL=71 (b). With increasing ETL, readout lobes and phase encoding blips are added on both sides of the spin echo, leading to a proportional increase in the ETL-duration. We maintained a constant k-space FOV (and image resolution) by varying the phase encoding gradient amplitude between the ETLs, similar to SENSE acceleration along the phase encoding direction. Each purple dot in the k-space trajectory represents a time step, and crosses indicate the gradient echoes (red to blue in temporal order). It is visible that gradient echoes occur at *K*_*x*_ = 0, indicating well-balanced readout gradient lobes and a proper choice of Δ*t*.

Each simulation used an ensemble of 10,000 protons. We used a set of identical random seeds for the resting and active state random walks. This seeding yielded the same set of diffusion paths in both physiological states, which allowed us to compute the BOLD effect at this ‘small’ number of protons; otherwise, a larger proton ensemble would be required to prevent slight differences in experienced encoding gradients from overshadowing the BOLD effect. Table C.3 gives an overview of the discretization parameters.

We simulated all odd ETL values in the realistic range of 27 to 77. Additionally, we included an ETL of 3 (too noise-corrupted in practice), which shows the theoretically achievable changes upon further minimizing the ETL-duration. We performed all simulations for two (ascending/pial) vein orientations: parallel to 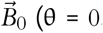, no vein-induced field inhomogeneities) and perpendicular to 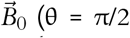, maximum vein-induced field inhomogeneities, as shown in Figure 1).

#### 2.3.2 Assumptions and simplifications

We made several assumptions and simplifications to create this biophysical model and apply it to SE-EPI simulations in GM and CSF.

First, we simplified the vascular morphology. We chose to represent vessels as infinite cylinders, which allowed us to use analytical field expressions. This simplification is defendable since vessel radii are generally much smaller than the corresponding lengths and curvatures.^21^We simulated microvessels with random orientations, which is a valid approximation as the capillary bed has very little preferential orientation and is relatively uniform.^9^Furthermore, we included *one large vein* per voxel, whereas multiple intracortical ascending and pial veins may exist in a GM and CSF voxel, respectively. Lastly, we neglected vessel-volume overlap. Since the goal is not to compare the simulations to experiments on a per voxel basis but to study the overall BOLD behavior across ETL-durations, we believe this approach suffices for our purpose.

Second, the diffusion of water protons was simplified. We considered only extravascular protons, as they constitute the dominant signal source at 7T^10^due to the short *T*_2_ and 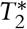 of blood at high field strengths.^3^ We did not allow any proton water exchange through the vessel wall, which is plausible given that the water exchange time between extra- and intravascular spaces is about 500 ms(*≫*TE=55 ms).^22^We also disregarded the inflow and outflow of protons into/out of the imaging voxel, as the extravascular diffusion paths are small compared to the voxel size.

Third, we simplified the susceptibility sources that generate the mesoscopic field. We assumed blood vessels of homogeneous susceptibility, while in reality the negative gradient of dHb-content along vessels is an additional source of field inhomogeneities and increases the BOLD effect. This effect is most notable in microvessels, as they are the primary location of oxygen extraction.^4^Additionally, we only considered field inhomogeneities induced by internal susceptibility sources. Even though the spin echo partially refocuses fields around large veins, we still expect the field inhomogeneities around large pial veins to be effective in GM voxels in reality.

Lastly, we used a few assumptions in the magnetization description. We simulated instantaneously applied pulses with a perfect flip angle, neglecting any *B*1-field inhomogeneities. Furthermore, we assumed instantaneously switching encoding gradients, whereas, in practical settings, the gradient fields are turned on/off at a slew rate. Considering the averaging effect of our finite integration time step, we do not expect considerable differences compared to implementing trapezoid gradients. Finally, we sampled only at the center of each readout lobe. This strategy will overestimate the k-space signal slightly as we do not account for 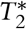 decay away from the gradient echo within the acquisition of a single k-line (*∽*1 ms). In reality, 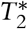 decay across the k-line will result in slight (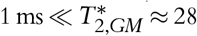 GM*≈* 28 ms) image-space blurring along the readout direction, which can reduce the precision of the measured BOLD values.^1^

### 2.4 In-vivo validation experiment

#### 2.4.1 Subject

We scanned one healthy subject (male, 26 years) in a supine position. Before the scanning, we obtained approval from our institution’s Medical Research Ethics Committee (MREC) and written informed consent from the subject. We executed the experiment according to the guidelines and regulations of the WMO (WetMedisch Wetenschappelijk Onderzoek).

#### 2.4.2 Scan protocol

The subject was scanned on a Philips 7T magnetic resonance imaging (MRI) scanner using a dual-transit head coil in combination with a 32-channel receive coil (Nova Medical, Wilmington MA, USA).

We acquired a *T*1-weighted MP2RAGE scan as structural reference.^23^The in-plane FOV was 230 × 230 mm^2^, and 160 slices of 1 mm thickness were acquired. We used an image matrix of 232 × 232, in-plane resolution of 1 mm, TE=2.2 ms, TR=6.2 ms (repetition time), TR_volume_=5500 ms, flip angle of 5°, and readout bandwidth of 405.1 Hz. The inversion times were 1000 and 3000 ms.

Precise targeting of end-tidal O2 partial pressure (PetO2) was accomplished using a RespirAct™system (Thornhill Research, Toronto, CAN). The RespirAct device consists of a computer-controlled gas blender. An output line connects the blender to a rebreathing circuit, taped to the subject to ensure an airtight seal using Tegaderm film (3 M, Maplewood, MN, USA). Two sensor lines are connected to monitor the end-tidal gas values and breathing pressure. The rebreathing circuit ensures that the subject inhales all delivered gas, facilitating precise targeting of end-tidal gas values.^24^

Each functional scan started with a 60 s baseline period (PetO_2_ = 100 mmHg), followed by a 180 s interval in which we targeted a PetO_2_ level of 650 mmHg, and reached effective levels of 600 mmHg. The hyperoxic stimulation thus yielded a ΔPetO2 of about 500 mmHg, which was high enough to observe a significant BOLD response. After activation, we programmed a 120 s recovery period for return to baseline. Figure 3 shows the target and effective PetO_2_ traces.

**Figure 3.**
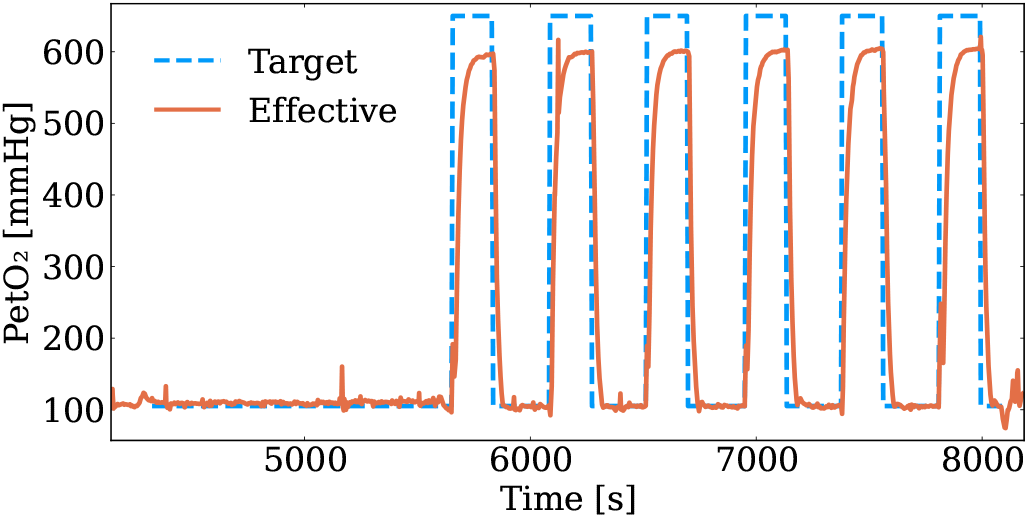
Target and effective end-tidal O_2_ pressures (PetO_2_) of the hyperoxic stimuli. The stimulus strength and duration were mostly consistent between the different SE-EPI acquisitions although we observed a peak in PetO_2_ in an early phase of the second stimulus (ETL=71 repeat scan). The CO_2_ pressures and the respiration rate during the stimuli are shown next to the PetO_2_ trace in Figure D.11.

We repeated the functional SE-EPI scans for five ETL (with implemented SENSE) values: 33 (3.8), 41 (3.1), 51 (2.5), 63 (2.0), and 71 (1.8). We randomized the ETL values to prevent any bias due to the potential history effects of the hyperoxic challenge. ETL=71 was the first scan, and we repeated this ETL to assess the repeatability of our experiment. The FOV was 227 mm in the readout- and 170 mm in the phase-encoding direction, and we acquired 30 slices. All scans used TE=55 ms, TR=4 s, and a symmetric readout window. Each SE-EPI scan consisted of 90 dynamics, preceded by three dummy cycles. We used SPIR to suppress fat signal contributions. Table D.1 presents the scan parameters (ETL, SENSE, k-space matrix, acquired resolution, water-fat shift, readout bandwidth) per ETL value. We preceded each SE-EPI scan with a corresponding top-up scan, consisting of a single dynamic (preceded by one dummy cycle) with identical parameters to the SE-EPI BOLD time series except for an opposite phase encoding gradient polarity. We used this scan to correct EPI distortions retrospectively.^25^The top-up scan and dummy cycles effectively extended the recovery period by 20 s to 140 s.

#### 2.4.3 Data pre-processing

We converted the raw DICOM data from the scanner to NIfTI format using the latest release of dcm2niix (v1.0.20230411).^26^Afterward, we performed a series of pre-processing steps using FSL functions,^27^as summarized in Figure D.4.

We extracted the brain based on the second inversion image of the MP2RAGE scan using FSL bet. Scanner software reconstructed a *T*_1_ map from the two MP2RAGE inversion images, based on which GM, WM, and CSF were segmented using FSL fast. We made a ventricle and cerebellum mask by registration of MNI152 atlases to *T*_1_

Each SE-EPI underwent subsequent motion correction (FSL mcflirt), top-up correction (FSL topup), and brain extraction (FSL bet). We registered the *T*1-space masks to the native space of each EPI using the boundary-based registration utility of FSL flirt. Registration of the GM, WM, and CSF segmentation resulted in three tissue-probability masks. We assigned the most likely tissue type to each voxel, allocating it to a single ROI. Lastly, we removed the top and bottom slices of the output images, as these EPI slices had artificially large intensity fluctuations due to motion-induced variations in FOV, leaving us with 28 slices in the output images. Each EPI was analyzed further in its own native space.

#### 2.4.4 Data analysis

For further analysis, we masked away the ventricles and cerebellum in the tissue-type ROIs to focus on cortical responses in GM and sulci-located CSF. Figure D.5 shows an example of the resulting GM and CSF segmentations.

We defined a baseline and activated period based on characteristic PetO_2_ time points that we logged during scanning. The differences between stimulus onset times between scans were maximally a single dynamic. Therefore, we used the same baseline and activated state intervals for each scan, using the most conservative intervals possible. We defined the baseline period as the interval before the stimulus was turned on, and the activated period as the interval after the peak PetO2 was reached but before the stimulus was turned off.

We performed temporal smoothing on a voxel-wise basis to remove high-frequency fluctuations, which do not reflect the BOLD response and are potentially associated with cardiac and respiratory variations. We used the rlowess function from MATLAB’s Curve Fitting Toolbox for this purpose, which performs a robust quadratic local regression.^28^We used a 20% smoothing window size (approximately the baseline duration) to reach a stable response during the activated period.

Finally, we calculated the BOLD maps by normalizing the SE-EPI signal to its voxel-wise average during the baseline period. We used the *median*, indicated by a superscript ‘*∽*’, in each tissue-type ROI to reflect the central tendency, which ensures robustness against outliers. Figure 4 shows an example of the BOLD maps and median %BOLD responses before and after temporal smoothing using rlowess.

**Figure 4.**
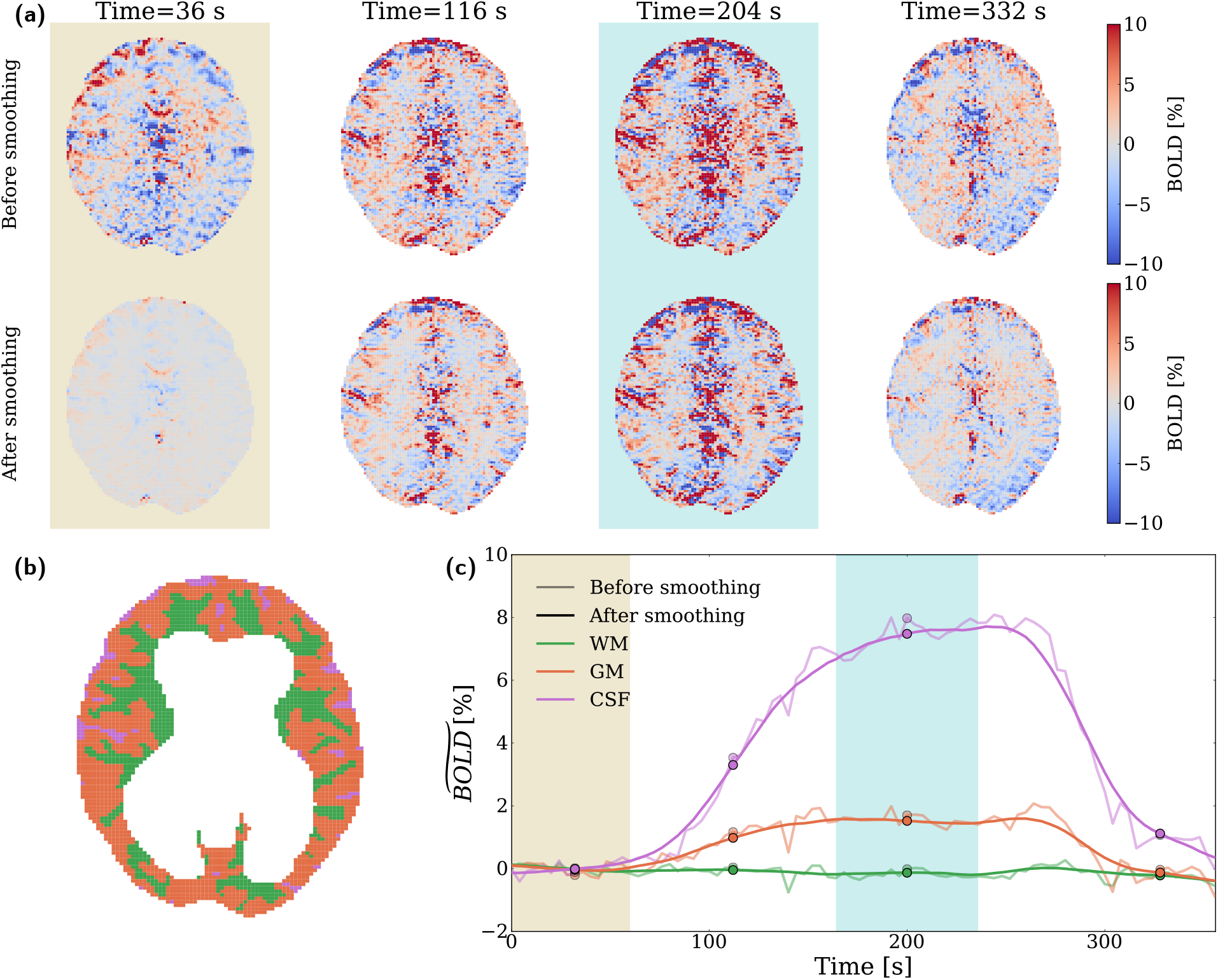
Representative BOLD maps and time traces (ETL=63), before and after temporal smoothing. The BOLD maps in panel (a) show a single transversal slice at four distinct time points: one in the baseline period (36 s), one during the build-up of the BOLD response (116 s), one in the activated period (204 s), and one during recovery to baseline (332 s). Panel (c) shows the median %BOLD responses across the full ROIs, where markers indicate the four selected time points. Yellow shading represents the baseline period, and cyan shading represents the activated period. Panel (b) shows the tissue-type ROIs in the transversal slice for reference. From panels (a) and (c), it can be seen that temporal smoothing using rlowess reduces not only temporal fluctuations but also increases spatial homogeneity. Notably, the baseline dynamic shows little to no BOLD response after smoothing. Besides the expected positive BOLD responses, many ROI voxels (mostly near the frontal lobe and in the brain interior) show a negative BOLD response.

## 3. Results

### 3.1 Simulation

Figure 5(a) shows the %BOLD signal change in the GM and CSF voxel as a function of ETL for the scenarios of a vein perpendicular and parallel to the *B*0 field.

**Figure 5.**
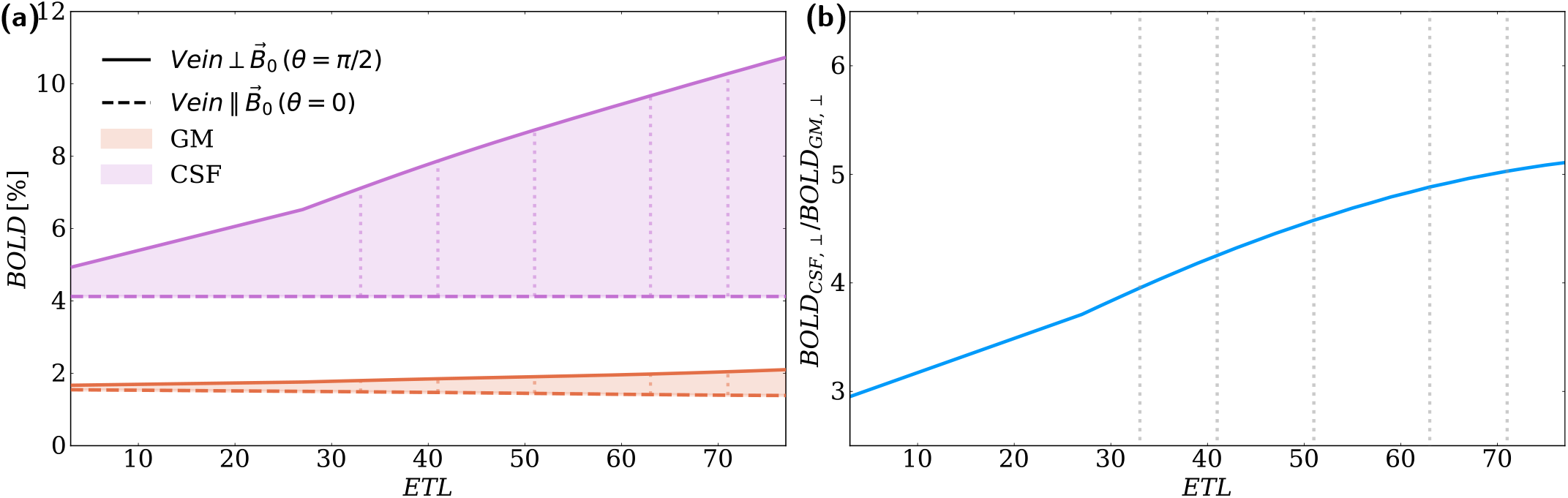
Simulated %BOLD signal changes in the CSF and GM voxel and their ratio as a function of ETL. The dotted vertical lines represent ETL values that we acquired in the validation experiment. Panel (a) shows the %BOLD signal changes for the scenarios of a vein perpendicular and parallel to the *B*_0_ field. In their perpendicular configuration, the veins induced maximum field inhomogeneities. The %BOLD increased as a function of ETL in both GM and CSF, in line with an expected increase in macrovessel contributions with increasing ETL (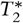 weighting). The absolute %BOLD increase as a function of ETL was larger in CSF than in GM, likely due to the large pial vein blood volume in CSF compared to the ascending vein blood volume in GM. Panel (b) shows the ratio of the CSF to GM %BOLD as our measure for macrovascular contamination in the perpendicular scenario. Our simulation predicted a consistent increase in the CSF to GM %BOLD ratio with increasing ETL (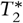 weighting). If we consider the range between the lowest (ETL=33) and the highest (ETL=71) ETL used in the validation experiment, we see a 30% increase. If we look at the range between ETL=3 and ETL =71, we see an increase of 70%. In their parallel configuration, the veins induced *no* field inhomogeneities. In the GM voxel, we dub this the ‘microvessel-only’ scenario and saw that the %BOLD reduced slightly as a function of ETL (panel a). In the CSF voxel, the %BOLD had an ETL-independent value of about 4% (panel a). Considering the absence of microvessels in the CSF voxel, there weren’t any vasculature-induced field inhomogeneities. Thus, the positive %BOLD must correspond to the simulated *T*_1_ shortening effect within the sulci-located CSF upon hyperoxic activation (Table C.1).

In their perpendicular configuration, the veins (one ascending vein in GM, one pial vein in CSF) induced maximum field inhomogeneities. The %BOLD increased as a function of ETL in both GM and CSF, in line with an expected increase in macrovessel contributions with increasing ETL (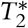 weighting). The absolute %BOLD increase as a function of ETL was larger in CSF than in GM, likely due to the large pial vein blood volume in CSF compared to the ascending vein blood volume in GM. Figure 5(b) shows the ratio of the CSF to GM %BOLD as our measure for macrovascular contamination in the perpendicular scenario. Our simulation predicted a consistent increase in the CSF to GM %BOLD ratio with increasing ETL (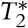 weighting). If we consider the range between the lowest (ETL=33) and the highest (ETL=71) ETL used in the validation experiment, we see a 30% increase. If we look at the range between ETL=3 and ETL =71, we see an increase of 70%. Note that an SE-EPI scan with ETL=3 would require a very high SENSE factor, which in practice would result in too much noise breakthrough.

In their parallel configuration, the veins induced *no* extravascular field inhomogeneities. In the GM voxel, we dub this the ‘microvessel-only’ scenario and saw that the %BOLD reduced slightly as a function of ETL (Figure 5, panel a). In the CSF voxel, the %BOLD had an ETL-independent value of about 4% (Figure 5, panel a). Considering the absence of microvessels in the CSF voxel, there weren’t any vasculature-induced field inhomogeneities. Thus, the positive %BOLD must correspond to the simulated *T*1 shortening effect within the sulci-located CSF upon hyperoxic activation (Table C.1). We confirmed that both the parallel and perpendicular CSF %BOLD curves shifted down by 4% when we used the same *T*1 value for the resting and active state of the simulation.

### 3.2 In-vivo validation experiment

#### 3.2.1 SEessed the quality of -EPI data inspection

We assour in-vivo SE-EPI data and will highlight our main findings; additional details and visualizations are provided in Appendix D.2. First, we found that the temporal SNR (tSNR) during the baseline period was in the same order of magnitude for the different scans (mean tSNR range = 36-47). Both ETL=71 scans scored the worst, which we expect to be caused by relatively large fluctuations in the respiration rate during these scans (Figure D.11). Second, the EPI images were structurally similar, and the registration quality to the *T*1 structural reference was consistent among scans. Third, we found the median to be a robust reflection of our data. Lastly, a linear regression between the two ETL=71 acquisitions indicated a bias between the scans.

#### 3.2.2 BOLD responses

Figure 6 shows the median %BOLD responses for each ETL acquisition in the GM and CSF ROIs. All acquisitions showed a distinct BOLD response in GM and CSF during the hyperoxic stimulus. The measured GM and CSF %BOLD responses in the activated state were in the same order of magnitude as our simulated %BOLD signal changes (Figure 5, panel a). The measured WM %BOLD responses (Figure D.10) were close to zero, which we expected considering the poor vascularization and low blood flow in this brain region.

**Figure 6.**
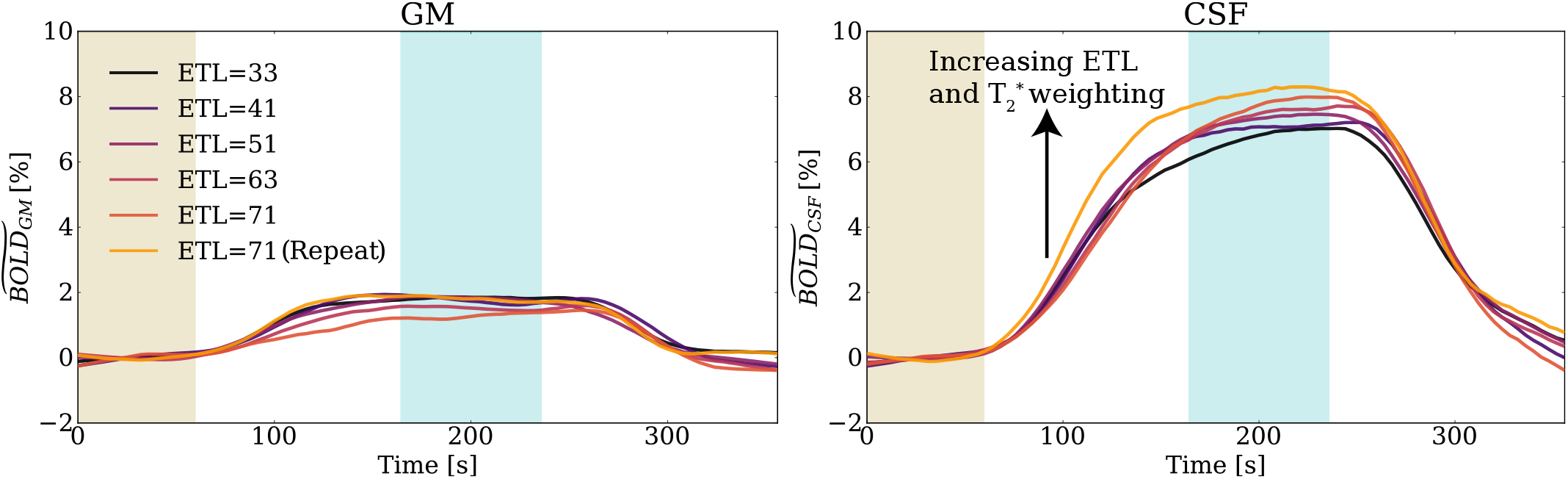
Measured median %BOLD responses for each ETL acquisition in the GM and CSF ROIs (WM %BOLD responses are also shown in Figure D.10). Yellow shading represents the baseline period, and cyan shading represents the activated period. All acquisitions showed a distinct BOLD response in GM and CSF during the hyperoxic stimulus. The measured GM and CSF %BOLD responses in the activated state were in the same order of magnitude as our simulated %BOLD signal changes (Figure 5, panel a). The GM %BOLD response tended to decrease as a function of ETL, whereas the CSF %BOLD response increased as a function of ETL. The consistent increase in the CSF %BOLD response as a function of ETL agrees with the simulated trend in Figure 5(a). Note that the ETL=71 repeat scan had an uncharacteristically high GM %BOLD response and a significantly different CSF %BOLD response than the other scans. For this reason, we focused on the first ETL=71 acquisition when we assessed the CSF to GM %BOLD ratio as a function of ETL.

We can see that the GM %BOLD response tended to decrease as a function of ETL, whereas the CSF %BOLD response increased as a function of ETL. The consistent increase in the CSF %BOLD response as a function of ETL agrees with the simulated trend in Figure 5(a). Note that the ETL=71 repeat scan had an uncharacteristically high GM %BOLD response (inconsistent with the trend across ETL) and a significantly different CSF %BOLD response than the other scans (mostly in the early phase). This is supported by the bias between the two ETL=71 acquisitions reported in Figure D.3. For this reason, we focused on the first ETL=71 acquisition when we assessed the CSF to GM %BOLD ratio as a function of ETL.

Figure 7 shows the computed CSF to GM %BOLD ratio across ETL. We included all time points in the activated state interval (for ratio over time, see Figure D.9); error bars reflect variations in the ratio across the activated state interval. Similar to the simulation (Figure 5, panel b), we observed an increase in the CSF to GM %BOLD ratio as a function of ETL (ignoring the ETL=71 repeat scan as discussed previously). The order of magnitude of the ratios agreed with our simulation. The computed increase in the CSF to GM %BOLD ratio across the range of ETL values (ETL range = 33-71) exceeded the simulated increase by a factor of two, at close to 60% increase.

**Figure 7.**
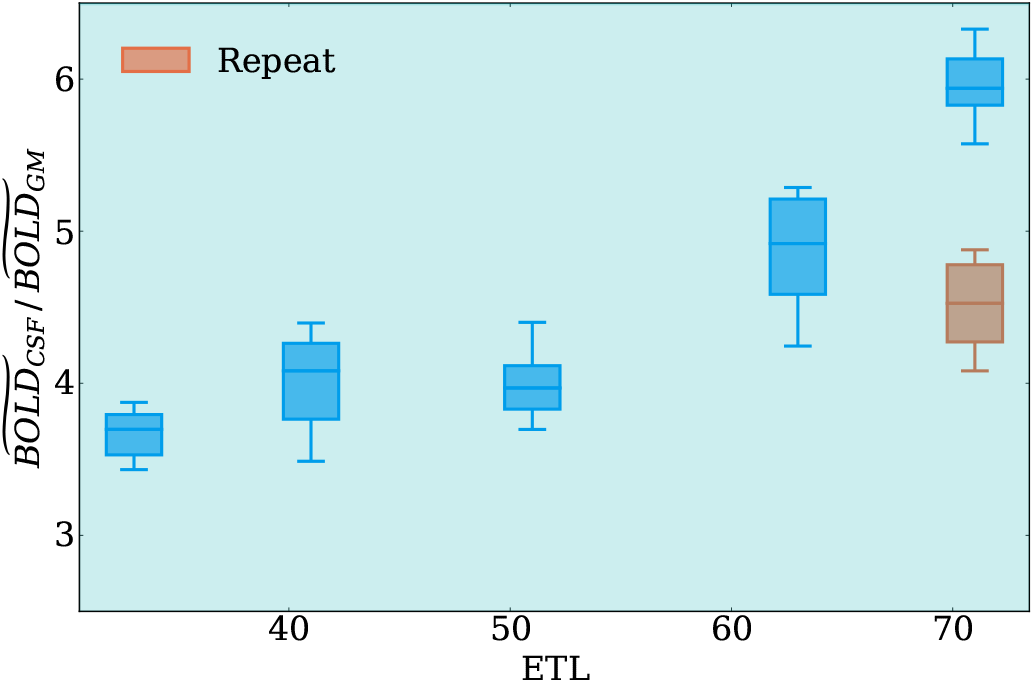
Computed CSF to GM %BOLD ratio across ETL. We included all time points in the activated state interval; the error bars reflect variations in the CSF to GM %BOLD ratio across the activated state interval. Similar to the simulation (Figure 5, panel b), we observed an increase in the CSF to GM %BOLD ratio as a function of ETL (ignoring the ETL=71 repeat scan as discussed previously). The order of magnitude of the ratios agreed with our simulation. The computed increase in the CSF to GM %BOLD ratio across the range of ETL values (ETL range = 33-71) exceeded the simulated increase by a factor of two, at close to 60% increase.

## 4. Discussion

The results of our study suggest that SE-EPI acquisitions, even though they use a refocusing pulse to reduce macrovascular contributions, still suffer from significant macrovascular contamination at realistic ETL values. The macrovascular contamination was apparent in our simulations by differences in the BOLD signals between parallel (‘microvessel-only’) and perpendicular large vein orientations. We probed the macrovascular contamination indirectly by computing the CSF to GM %BOLD signal ratio, which we also measured in a validation experiment at multiple ETL values. Both our simulation and validation experiment showed an increasing probed macrovascular contamination with increasing ETL and thus 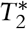 weighting. This trend agreed with the increase in differences between the parallel and perpendicular simulated BOLD signals as a function of ETL. In our simulation, we predicted a 30% macrovascular contamination increase between ETL=33 and ETL=71. In our validation experiment, we computed a difference of about 60% between these ETL values.

The steep increase in macrovascular contamination as a function of ETL is explainable by an asymmetric SE (ASE) simulation of the BOLD signal. Analogous to a SE-EPI experiment, asymmetric (relative to the SE) sampling moments increase 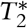 and hence macrovascular weighting. Similar to the work of Boxerman et al., we employed an ‘infinite’ voxel model filled to a fixed volume fraction with cylinders (vessels) of a uniform radius in random orientations. We performed a sweep of the vessel radius ranging from 1 to 100 μm, covering both micro- and macrovessel radii, and plotted the dependence of the BOLD signal on the vessel radius and the sampling moment. The ASE simulation (Figure B.1) shows that at the SE time point (similar to an ETL that approaches 1), the signal contributions of large veins will be close to zero, and the SE BOLD signal is most specific to radii in the microvascular range. Further from the SE time point, the sensitivity to large veins steeply increases and causes the overall microvascular specificity to decrease. With increasing ETL-duration in a SE-EPI experiment, we include more asymmetric sampling moments, which leads to an increase in macrovascular contamination and an associated decrease in microvascular specificity.

In the validation experiment, we observed a decrease in the GM BOLD response with increasing ETL. This decrease was also seen for the simulated parallel ‘microvessel-only’ scenario and hints at prevailing microvascular contributions in the measured GM BOLD response. This decrease disappeared in our simulation upon removing the encoding gradients, which suggests that the cause is encoding gradient-induced dephasing. By design, the readout gradient refocuses stationary protons at each gradient echo. Proton diffusion will result in a loss of refocusing and a signal reduction. With an increasing ETL, the readout window becomes longer, and we expect more gradient-induced signal loss.

Multiple limitations in our simulation can explain differences between the simulated and measured BOLD responses and macrovascular contamination.

Most importantly, we severely simplified the vascular morphology to a single collection of infinite cylinders, even though we took care to choose our vascular morphology parameters in the GM and CSF voxel according to available literature describing the human^29,30^and macaque^2^brain. The human brain microvasculature has relatively well-described statistical properties and resembles a dense network of randomly oriented vessels, which we expect our model captures appropriately.^30^Much less is known about the (variable) morphology of sparsely distributed intracortical ascending veins in GM and pial veins in CSF. We simulated a single vein in both GM and CSF but observed that our simulated measure for macrovascular contamination depends heavily on the orientation, radius, and number of large veins. Recently, several studies have been making an effort to create statistical models that approximate vascular morphologies of rodents^31^and humans.^32^To closer resemble the BOLD contrast and its specificity in the human brain, these studies could serve as a starting point for a switch to a more realistic and complicated representation of the vascular network. Multiple vascular networks can be employed to predict spatial variations in the BOLD specificity.

Furthermore, we only simulated internal (to the voxel) susceptibility sources in this study. Realistically, pial veins generate far-stretching field inhomogeneities, which are a source of macrovascular contamination even outside the voxel that contains them. The effects of external susceptibility sources will increase with ETL-duration due to an increase in *T*\* weighting, which most strongly increases undesired large vein contributions in the GM BOLD signal. Consequentially, provided a realistically simulated vascular morphology, our simulated CSF to GM ratio and its increase as a function of ETL-duration might exceed the measured values. External susceptibility sources could be included to make the simulation more realistic, effectively simulating a collection of neighboring voxels. Since large vessels have the most far-reaching effects, it could be sufficient to account only for large pial veins.

During our validation experiment, we came across some challenges and limitations. Additionally, some analysis steps could be added or refined in the future.

The first challenge in our type of experiment is the sensitivity to partial volume effects (PVEs). Whereas our simulations focused on a pure GM and pure CSF voxel, this was not feasible in our experiment due to the finite voxel size. As a part of data pre-processing, we assigned each voxel the most probable tissue type. As a result, many voxels in our experimental ROIs will be a combination of multiple tissue types. We tried using a minimum probability threshold to alleviate PVEs, but this drastically reduced the number of CSF voxels in our analysis. With a one-subject sample size, this was not a feasible solution. Scanning multiple subjects could allow the selection of smaller ROIs with higher confidence for corresponding to GM or CSF. Additionally, using different scanner hardware or sequence designs could enable a reduction in resolution and ROI definitions with fewer partial volume voxels.

Second, some slight inconsistencies in ROI definitions across ETL might have hindered our comparison. We analyzed each EPI in its own native space, and its exact ROIs depended on the EPI itself (including its spatial discretization and potential artifacts) and the registration quality from *T*1 to EPI space. Although the quality of the EPI data and registrations seemed to be consistent across ETL (see Appendix D.2), we still recommend analyzing all SE-EPIs in a common image space in the future to ensure ROI consistency.

Third, the dynamic BOLD behavior of the ETL=71 repeat scan deviated significantly from the other acquisitions, and we saw a bias between the BOLD activation patterns of the ETL=71 scans. The tSNR and motion (correction) parameters were highly similar between the two ETL=71 scans and are unlikely to have caused these differences. We observed a rapid peak in PetO2 and a dip in PetCO2 in an early phase of the repeat scan’s hyperoxic stimulus (Figure D.11). As the BOLD response is highly sensitive to small absolute signal changes and represents a cumulative change, a short physiological disturbance could translate significantly into the measured BOLD response. It is unclear what caused this disturbance. An additional scan session is needed to re-assess the repeatability of our experiment for multiple subjects.

Lastly, it would be valuable to consider only voxels that respond significantly to the hyperoxic stimulus. Even though we assumed that the administered hyperoxia affects the whole brain relatively homogeneously, this will not hold perfectly in reality. Instead of selecting ROIs solely based on tissue type and spatial location (i.e. not in the ventricles or cerebellum), we could also account for dynamic behavior. We could achieve this by employing a general linear model to select only significantly responding voxels based on their p-value.

Finally, we would like to reflect on our choice of hyperoxia as a proxy for functional stimulation. Although the prospects of a global dHb change with presumably minimal CBV changes motivated us to use hyperoxia, we experienced some drawbacks in practice.

At our 600 mmHg PetO2 target, significant *T*1 shortening is expected in the CSF ROI and was accounted for in our simulations. *T*1 shortening occurs under hyperoxic conditions due to the inflow of excess paramagnetic molecular O2 into the CSF; it forms an additional source of positive signal change upon activation, next to the decreased dHb level. In our simulation, we implemented the *T*1 shortening effect based on *T*_1_ measurements of CSF in the sulci at variable PetO_2_,^24^giving a contribution of *∽*4% to the BOLD signal (constant across ETL). As our measure for macrovascular contamination reflects *relative* changes in the CSF versus GM BOLD level, the *T*1 shortening affects the resulting CSF/GM BOLD trend across ETL. In our simulation, we found that this shortening effect significantly alters the ratio’s trend compared to an otherwise equivalent simulation without *T*1 changes. Consequentially, the *T*1 shortening mixes non-dHb related effects in our measure for macrovascular contamination and makes our simulation depend heavily on the correctness of the *T*1 values. A strategy to experimentally mitigate the contribution of *T*1 shortening could be to fit *T*1 in a multi-echo SE-EPI experiment and retrospectively correct for *T*1 shortening.

Additionally, we observed prevalent *negative* BOLD responses (35-43% of all voxels in the GM ROI, 11-21% of all voxels in the CSF ROI). Figure D.8 shows the BOLD responses in voxels with an overall positive and negative BOLD contrast. Although this observation opposes our initial expectation of a hyperoxia-induced global *positive* BOLD contrast, some potentially underlying mechanisms have been reported in literature.^14,33^Pilkinton et al. suggested that paramagnetic O2 gas in the upper airway results in increased *B*0 inhomogeneities (and decreased 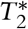 values in surrounding regions) in the active state, consequentially reducing the signal and increasing distortions.^14^In line with this hypothesis, we observed significant negative BOLD contrast near the frontal lobe and along the brain periphery in the phase encoding direction. Song et al. attributed negative BOLD contrast to an increase of molecular O2 concentrations and resulting field inhomogeneities (and 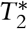 reduction) *in the extravascular space*, supported by blood gas and NMR analysis under hyperoxic conditions.^33^Opposed to the work of Pilkinton et al., Song et al. observed negative BOLD contrast primarily in the subcortical area. In our work, we observed negative BOLD contrast in both the brain periphery and the subcortical region. Furthermore, we observed an increased prevalence of negative BOLD contrast with increasing ETL (and 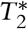 weighting), in line with both mechanisms. Lastly, vasoconstriction could contribute to negative BOLD responses. Whereas some sources describe hyperoxia as non vaso-active,^14,34^vasoconstriction has been reported in several other studies that typically used long stimuli compared to our study (*∽*10-30 min versus 2 min).^33,35^ The extent of vasoconstriction likely depends on the stimulus strength and duration.

Thus, although both functional and hyperoxic activation reduces local dHb concentrations, some key differences might hinder the transferability of our results to functional MRI. During functional activation, the main driver of dHb decreases is an increase in blood flow (and oxygen supply); during hyperoxic activation, it is a preferential metabolism of abundant plasma-dissolved oxygen over hemoglobin-bound oxygen. Contrary to functional activation, *non*-negligible paramagnetic effects of molecular oxygen accompany hyperoxic activation. In our study, these molecular effects likely contributed towards *T*1 shortening and negative BOLD contrast. To alleviate the molecular O2 contributions, we could target lower PetO2 levels at the cost of smaller absolute BOLD responses. Alternatively, we might consider a more direct approach using a *functional* stimulus, such as a visual checkerboard pattern stimulus.^3^In the latter case, CBV changes will become significant and should be accounted for in our model, increasing the complexity of the simulation.

Several studies have previously characterized macrovascular contamination of the BOLD signal based on laminar fMRI of the visual cortex using *functional* stimuli.^1,3,36^To probe the microvascular specificity, these studies used knowledge of the regional vascularization of the tissue across lamina; superficial GM is adjacent to the vein-rich pial surface, whereas deeper GM consists of microvessels with a sparse distribution of ascending veins. Ergo, a decrease in the BOLD signal in superficial GM under constant deep GM signal indicates a reduction of macrovascular contamination.^1,3^

Goense et al. studied the dependence of the macrovascular contamination on the SE-EPI readout duration in the macaque brain at 4.7T using multi-shot SE-EPI with a variable number of segments.^3^They found that reducing the SE-EPI readout duration (and 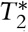 weighting) decreases the superficial GM BOLD signal, whereas the activation in deep GM was relatively insensitive. Correspondingly, we saw a simulated and experimental decrease in macrovascular contributions with decreasing ETL-duration and a relatively constant simulated ‘microvessel-only’ GM BOLD signal. Goense et al. suggested a dominance of microvessel contributions for readout durations below 20 ms 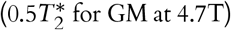. By extension, this would correspond to about 14 ms (ETL=20) in our 7T study. In agreement, our simulation shows a comparable BOLD signal in GM and CSF (excluding *T*1 shortening) at ETL=20.

Pfaffenrot et al. used a *T*2-prepared sequence with short multi-echo GE readouts to study the macrovascular contamination as a function of the GE echo time (*TE*_*GE*_) in humans at 7T.^1^They observed a prominent peak at the pial surface in their data (representing macrovascular contamination) that reduced with decreasing *TE*_*GE*_(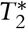 weighting). Laminar specificity was dramatically affected by 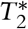 weighting even for *TE*_*GE*_ as short as 2.25 ms, which Pfaffenrot et al. largely attributed to GM-CSF PVEs at the pial surface. In accordance, our simulation study indicates that even at ETL=3 (2.1 ms ETL-duration), the CSF BOLD signal (excluding *T*1 shortening) is close to 1%, which is significant compared to the *∽*1.7% GM BOLD signal.

Alternative to reducing the ETL-duration, focusing on the early phases of the BOLD response could also help to increase the microvascular specificity. Siero et al. used the temporal characteristics of the SE-EPI and GE-EPI hemodynamic response functions (HRFs) to probe microvascular specificity in humans at 7T.^36^A high similarity between the SE-EPI (relatively insensitive to macrovascular contamination) and GE-EPI HRFs in early phases of the BOLD response indicated that early phases of the HRF are weighted towards the microvasculature.

## 5. Conclusion

Using SE-EPI acquisitions does not guarantee spatial specificity to the microvascular component of the fMRI BOLD signal in humans at 7T. However, careful sequence design choices may help to reduce macrovascular contamination as much as possible. In this work, we developed a biophysical model that was used to assess macrovascular contamination and is easily extendable to numerous other use cases in terms of vascular organization, magnetic field strengths, and pulse sequences. Whereas we only simulated ‘static’ BOLD responses (the resting and active physiological state) in this work, we could use a time-dependent oxygenation level to predict dynamic BOLD responses in the future.

To reduce macrovascular contamination, it is beneficial to use an ETL-duration that is as short as possible and to minimize PVEs as much as possible. The SE-EPI BOLD signal is more sensitive to pial veins in CSF than to the targeted GM microvasculature, especially at long ETL-durations (>14 ms). Reducing the ETL-duration from 49.7 ms (ETL=71) to 23.1 ms (ETL=33) decreases macrovascular contamination as probed by the CSF to GM %BOLD ratio by up to 30% according to our simulation, and by up to 60% according to our validation experiment. The reduction of the ETL-duration through SENSE acceleration is limited by noise breakthrough upon increased k-space undersampling. Provided this limit, it is additionally crucial to prevent GM-CSF PVEs that magnify the GM BOLD signal, heavily reducing its microvascular specificity.

To improve the value of SE-EPI BOLD acquisitions, we propose an alternative strategy to reduce the ETL-duration and the associated macrovascular contamination. Once the SENSE acceleration has reached its maximum potential, the ETL-duration can be shortened further by reducing the echo spacing (set by the readout gradient lobe duration). Recently, more advanced gradient insert coils are becoming available that achieve higher gradient amplitudes^37^and can thus cover the same k-space FOV faster, without any additional undersampling penalty. It will be interesting to see how much gain in microvascular specificity such a gradient system will achieve.

## 6. Acknowledgements

We would like to express our gratitude to Wouter Schellekens (Donders Institute, Radboud University, NL) for helping to set up the gas-challenge experiment. This work was supported by the National Institute of Mental Health of the National Institutes of Health under the Award Number R01MH111417 and the Dutch Research Council under the Award Number 18969. The content is solely the responsibility of the authors and does not necessarily represent the official views of the National Institutes of Health.

## 7. Code Availability Statement

The code that supports the findings of this study is openly available from the corresponding author.

## Appendices

### A Bloch matrices

We integrate the magnetization of each proton through a sequence of matrix multiplications. The magnetization 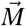 is initialized at steady-state according to:

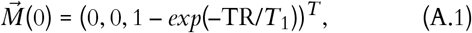

where TR is the repetition time of the pulse sequence. At each time point *t*, the magnetization at the next time point, *t* + Δ*t*, is calculated as:

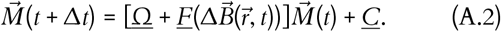

Here 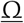 represents instantaneous pulse action (rotation of the magnetization vector) in a left-handed coordinate system with azimuthal angle α and polar angle β dictating the pulse orientation:

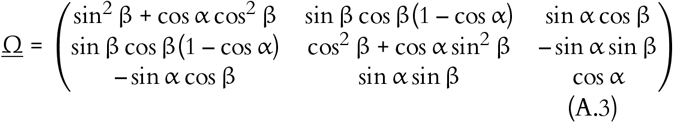

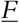 and 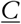 comprise NMR relaxation and precession at a field offset 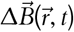:

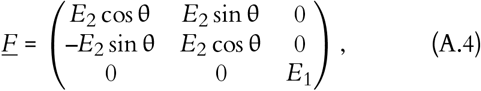

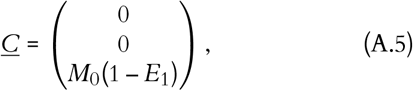

with an incremental phase change 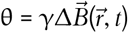, longitudinal relaxation *E*_1_= exp –Δ*t*/*T*_1_ and transversal relaxation *E*_2_= exp –Δ*t*/*T*2. γ is the gyromagnetic ratio of hydrogen, equal to 2π·42.58 MHz/T. <u>C</u> reflects the recovery of the longitudinal magnetization to its thermal equilibrium value *M*0.

### B. ASE simulation

An ‘infinite’ (size proportional to the vessel radius) voxel was filled with 800 cylinders of uniform radius in random orientations. A 10,000 proton ensemble underwent a random walk in the extravascular space (Δ*t*=0.1 ms). A sweep of the radius was performed to investigate the dependence of the %BOLD signal on the vessel radius and the sampling moment; τ is the temporal offset to the SE that defines the sampling moment. The ASE simulation parameters were matched to the SE-EPI simulation parameters for microvessels in GM (as described in Section 2.2). We did not account for variations in Hct and *Y* that would normally exist between microvessels and veins and no spatial encoding was applied. Figure B.1 shows the dependence of the ASE %BOLD signal on the vessel radius and sampling moment.

**Figure B.1.**
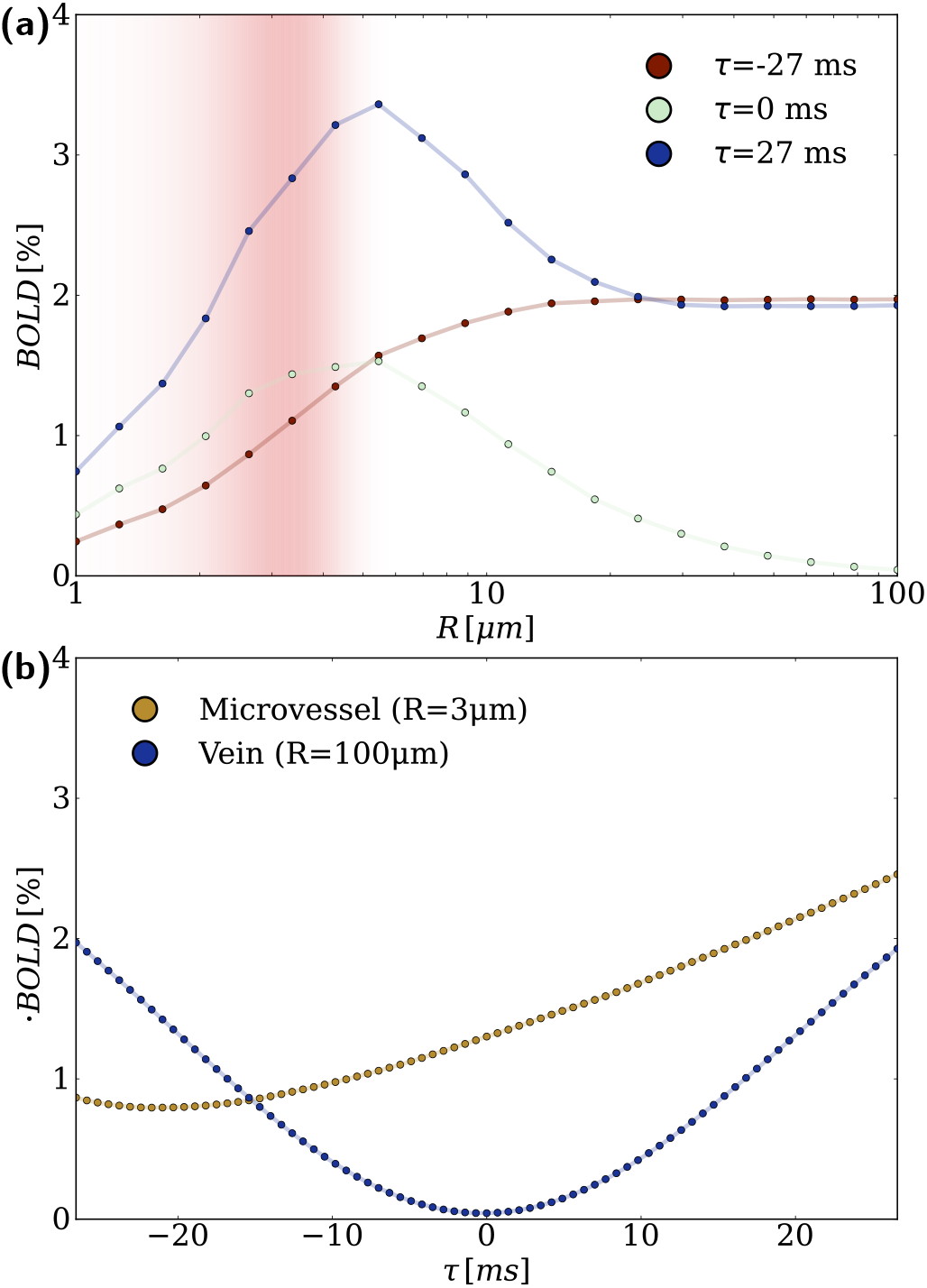
Dependence of the ASE %BOLD signal on the vessel radius and sampling moment. In panel (a), the dependence of the BOLD signal on the vessel radius is illustrated. In red, we indicated the microvessel radius distribution used in this paper. Clearly, we are primarily sensitive to microvessels at the SE (τ=0). At large offsets to the SE, the microvascular specificity is much lower, mostly due to an increased sensitivity to macrovessels. In panel (b), the dependence on τ is illustrated. Each marker corresponds to the timing of a gradient echo in the analogous SE-EPI scenario. As τ increases (further from the SE), we become more sensitive to large veins as we move further from the refocusing point and 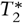 weighting increases. The sensitivity to microvessels generally increases over time as the phase dispersion grows when protons spend more time in the inhomogeneous mesoscopic field.

### C SE-EPI simulation

**Table C.1.**
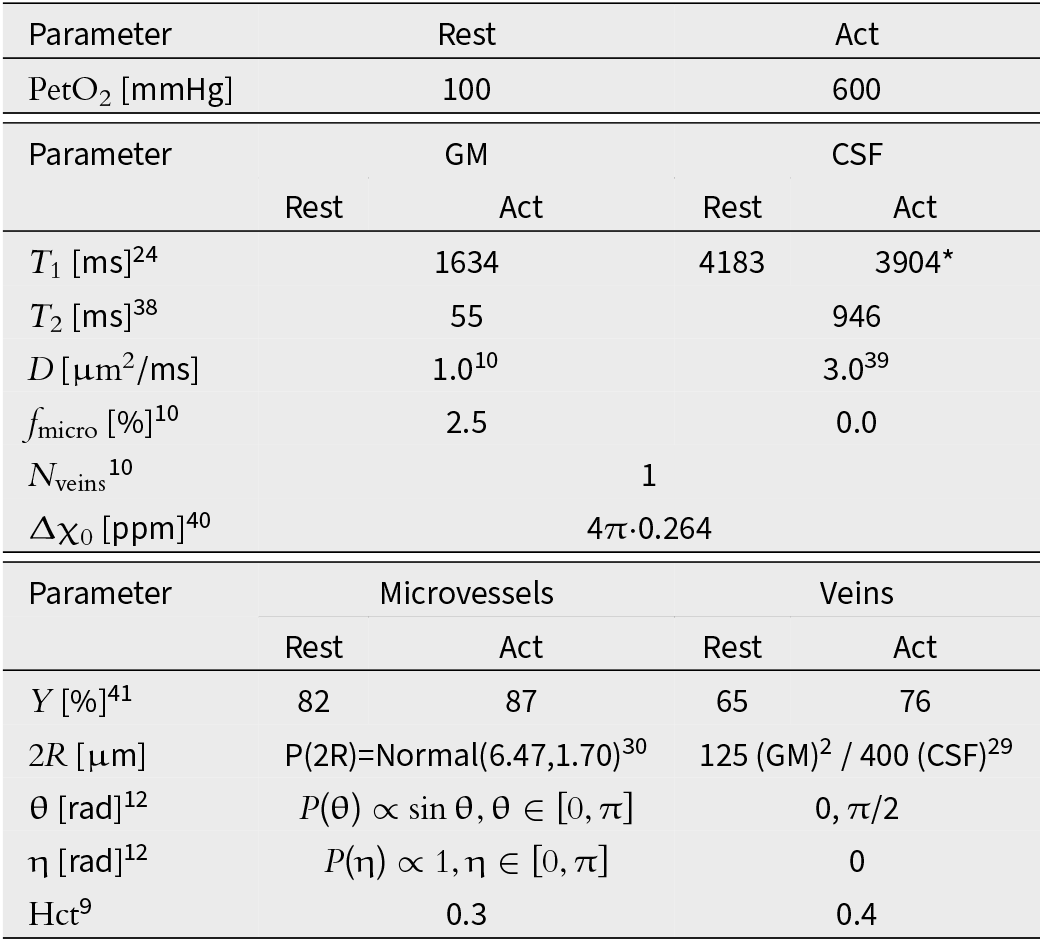
Physiological simulation parameters. We obtained the *T*_1_-value denoted with (*) from the relaxivity fit for *T*_1_ in the sulci described in [24] at an end-tidal O_2_ partial pressure (PetO_2_) level of 600 mmHg (Figure 3). We simplified the macrovasculature to a single vein in both GM and CSF.

**Table C.2.** Pulse sequence simulation parameters. We only simulated odd ETL values, which include the acquisition of the central k-space line.

| Sequence parameter | Value |
| --- | --- |
| Main field strength ($B_0$) [T] | 7 |
| Echo time (TE) [ms] | 55 |
| Echo spacing [ms] | 0.7 |
| Repetition time (TR) [ms] | 4000 |
| Isotropic resolution ($\Delta x_{iso}$) [mm] | 1.8 |
| Phase encoding blip duration [ms] | 0.15 |
| ETL values | 3, 27-77 (odd) |

**Table C.3.**
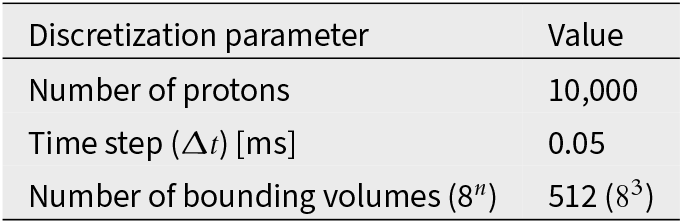
Discretization simulation parameters. The discretization time step applies to both the random walk and pulse sequence.

### D In-vivo validation experiment

#### D.1 SE-EPI acquisition parameters

**Table D.1.** Scan settings that differed between the SE-EPI acquisitions. In developing the protocol, we ensured that the k-space matrix size (x by y), acquired resolution (Δ*x*_acq_, x by y by z), and readout bandwidth (BW_read_) were approximately constant between scans. We did this to allow for a fair comparison to the simulations and to prevent large differences in (t)SNR between the scans. The water-fat shift (WFS) in pixels increases with increasing ETL due to a decreased phase encoding bandwidth.

| ETL / SENSE | Matrix | $\Delta x_{acq}$ [mm<sup>3</sup>] | WFS [pix] | $BW_{read}$ [Hz] |
| --- | --- | --- | --- | --- |
| 33 / 3.8 | 96x124 | 1.77x1.82x1.80 | 24.26 | 2010.4 |
| 41 / 3.1 | 96x124 | 1.77x1.81x1.80 | 29.97 | 2010.0 |
| 51 / 2.5 | 96x125 | 1.77x1.81x1.80 | 37.11 | 2009.7 |
| 63 / 2.0 | 96x126 | 1.77x1.80x1.80 | 45.67 | 2010.1 |
| 71 / 1.8 | 96x126 | 1.77x1.80x1.80 | 51.37 | 2010.8 |

#### D.2 SE-EPI data quality

We assessed the quality of our in-vivo SE-EPI data after the series of pre-processing steps described in Section 2.4.3 and visualized in Figure D.4. First, we characterized the temporal SNR (tSNR) during the baseline period. The resulting tSNR maps are shown in Figure D.1, and whole-brain mean and standard deviations are reported in Table D.2. The mean tSNR values in the brain were in the same order of magnitude (tSNR range = 36-47) for the different scans, with both ETL=71 scans scoring the worst. We expect that the relatively large fluctuations in respiration rate during the ETL=71 scans (see Figure D.11) caused the lower tSNR values of these scans. Second, a visual assessment showed a good alignment between the *T*1 map and its segmentations to the EPI data (see Figure D.2). Third, we computed the cross-correlation (CC) between each EPI and the *T*1 map (in EPI space). The CC scores were all 0.60 or 0.61, showing that the registration quality was consistent. Lastly, we verified the structural similarity between the EPI images. We took ETL=63 as the reference, as it had the highest tSNR, and registered all other EPIs to its native space. The CC scores to the reference EPI all fell in the range of 0.94-0.98, verifying that the scans were structurally consistent and ETL-dependent distortion corrections were successful. Table D.2 reports all CC-scores.

Next, we inspected the BOLD values resulting from the data analysis steps in Section 2.4.4. Figure D.6 shows that the distributions of BOLD values per ROI are approximately Gaussian (outliers exist far from the defined bins). We thus expect the median to reflect the average BOLD signal in an ‘average’ voxel well. Figure D.7 compares the mean and median measures of central tendency and illustrates that the median is much less reliant on the temporal smoothing; additionally, we visually compared rlowess smoothing to a wavelet smoothing approach and observed that rlowess filters out high-frequency fluctuations more effectively.

To determine the repeatability of our experiment, we assessed the spatial similarity of the activated state BOLD maps for the ETL=71 scans. Figure D.3 shows the density scatterplot of the ETL=71 scan 1 versus scan 2 (repeat) BOLD comparison within the GM ROI. A bias is apparent from the nonzero slope (0.72) and offset (0.59) of the linear regression (*R*^2^=0.37).

**Figure D.1.**
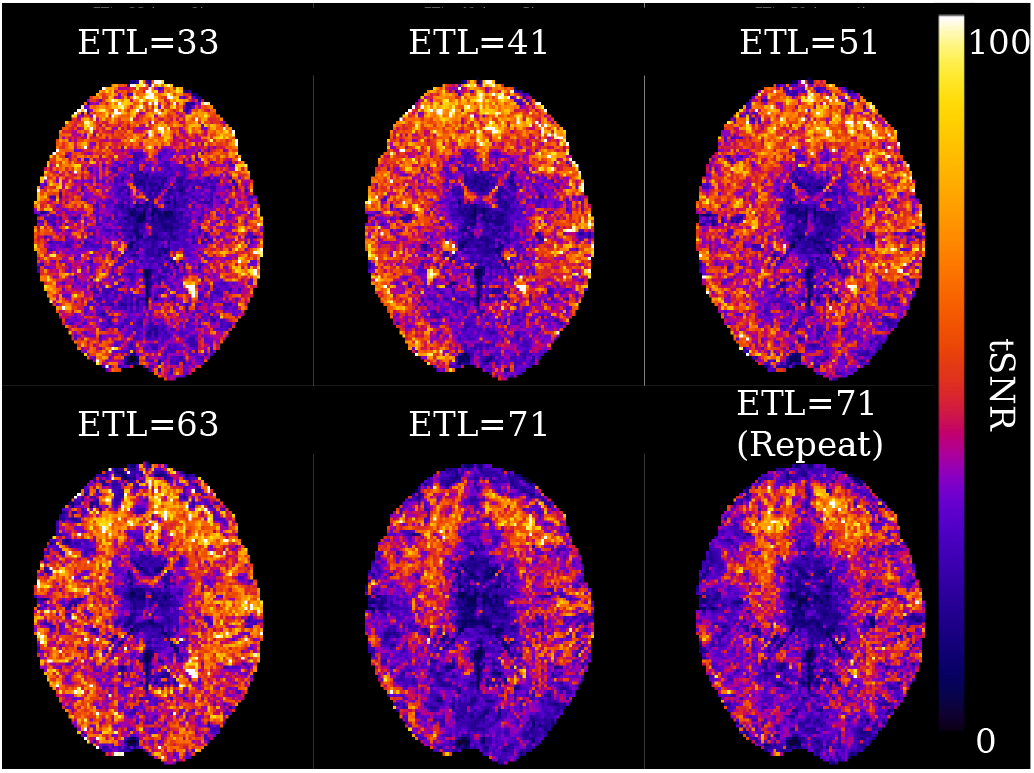
tSNR maps for all SE-EPI BOLD scans at a similar transversal location. We selected the baseline dynamics and divided the temporal mean by the standard deviation to compute the tSNR maps. Even though the absolute values are in the same order of magnitude, the ETL=71 scans have a notably lower overall tSNR than the other scans. We hypothesize that the differences in tSNR between the scans relate to respiratory stability; the respiration rate plotted in Figure D.11 shows relatively high fluctuations during the first two (ETL=71) scans.

**Table D.2.** Quality measures for each scan. Scan # indicates the order in which the scans were acquired. μ_tSNR_ and Book Antiqua σ_tSNR_ are the whole-brain tSNR mean and standard deviation, respectively. Cross-correlation scores were computed within the brain. The EPI with the highest tSNR (*) was taken as the reference for comparing the EPIs amongst each other through cross-correlation (CC).

| ETL / SENSE | Scan # | $\mu_{\text{tSNR}}$ | $\sigma_{\text{tSNR}}$ | CC to T1 | CC to EPI* |
| --- | --- | --- | --- | --- | --- |
| 33 / 3.8 | 6 | 43 | 19 | 0.60 | 0.96 |
| 41 / 3.1 | 5 | 45 | 19 | 0.60 | 0.98 |
| 51 / 2.5 | 4 | 44 | 18 | 0.60 | 0.98 |
| 63 / 2.0* | 3 | 47 | 20 | 0.60 | 1.00 |
| 71 / 1.8 | 1 | 38 | 16 | 0.60 | 0.96 |
| 71 / 1.8 | 2 | 36 | 16 | 0.61 | 0.94 |

**Figure D.2.**
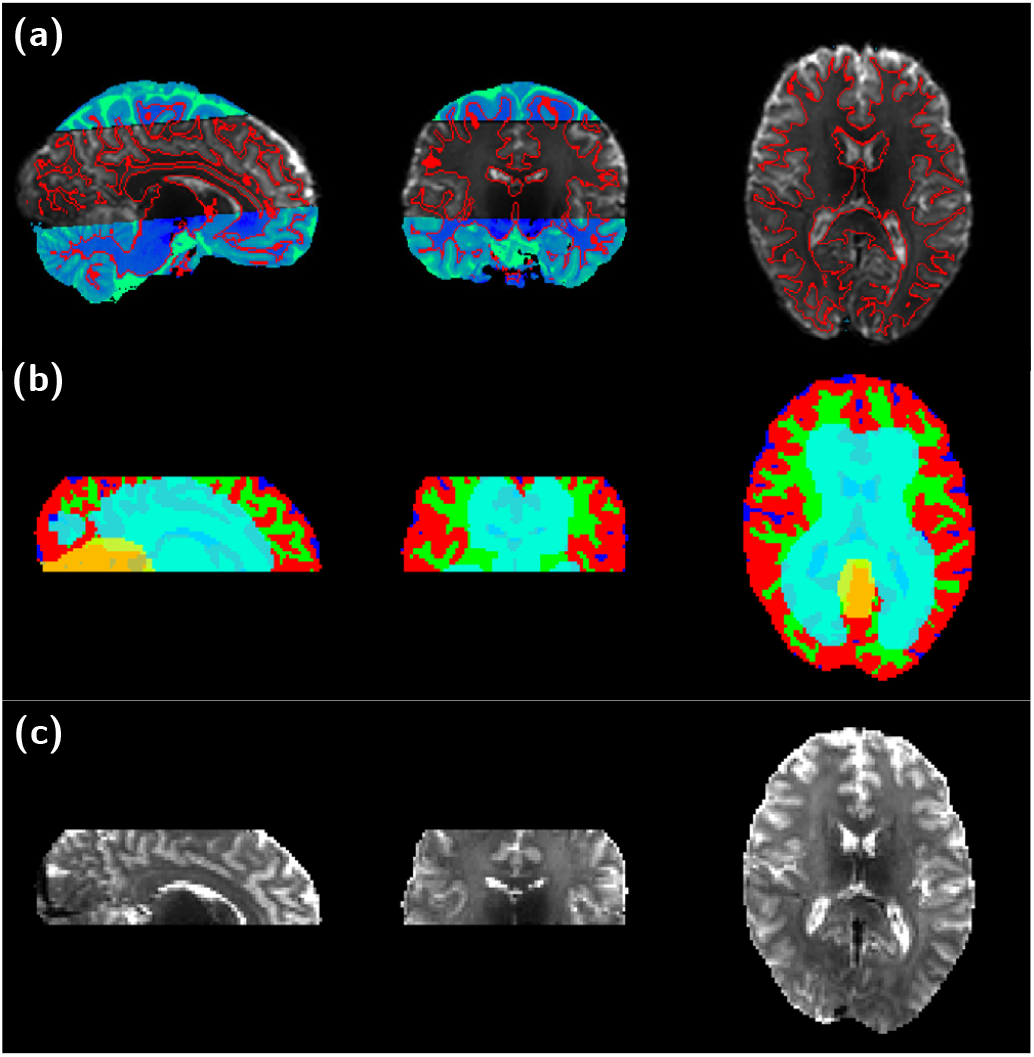
Visual assessment of the alignment between the *T*_1_ map and its segmentations to a SE-EPI intensity image (ETL=63). Panel (a) shows the alignment in *T*_1_ space. The EPI (grayscale) was registered to *T*_1_ space and overlaid on the *T*_1_ map (blue-green). The WM boundary indicated in red was used for the registration, maximizing the EPI intensity difference across the boundary. The red boundary overlaps nicely with the visible contrast change from WM to GM/CSF in the EPI, and the *T*_1_ and EPI map are well-matched at the EPI FOV boundary. Panel (b) and (c) together show the alignment in EPI space. In panel (b), green, red, and blue represent the respective WM, GM, and CSF segmentations. Cyan represents the ventricles, and yellow represents the cerebellum. Panel (c) shows the EPI intensity image at the same orthogonal slices. The masks correspond well with the visible contrast and structures in the EPI intensity image.

**Figure D.3.**
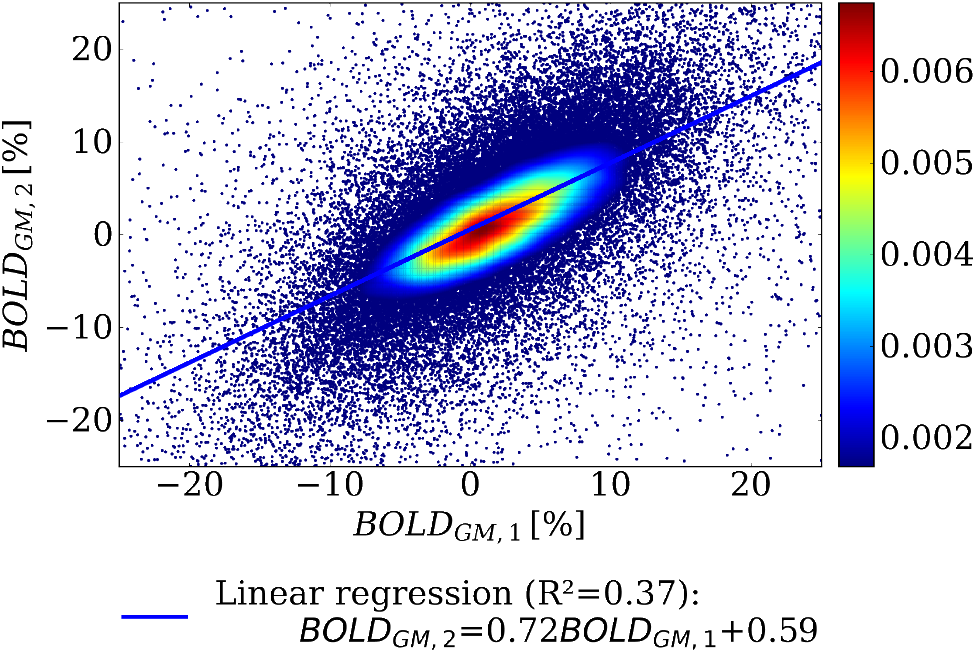
Scatterplot of the activated state GM BOLD values in the first versus second acquisition with the ETL=71 scan protocol. The BOLD maps were averaged across the activation period and values outside of the range of -25 to 25 were not taken into account. A linear regression was performed for a voxel-wise comparison of the BOLD values. The linear regression (*R*^2^=0.37) indicates a bias between the BOLD values of the two scans, with a slope of 0.72 and an offset of 0.59. We expect that this bias is related to observed fluctuations in the end-tidal gas levels during the acquisition of scan 2, see Figure D.11.

#### D.3 Pre-processing pipeline

**Figure D.4.**
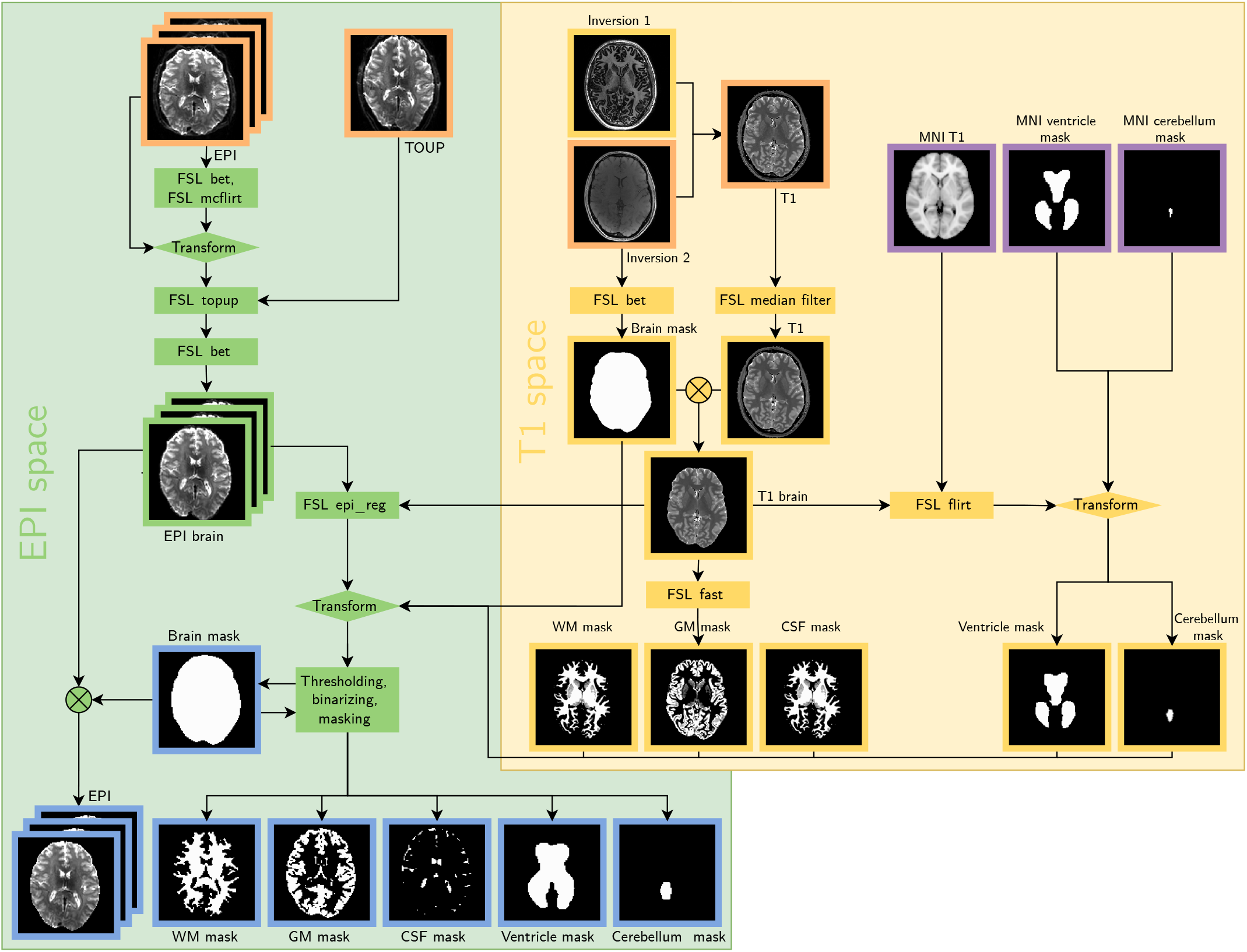
The pre-processing pipeline, implemented using FSL.^27^Orange outlines indicate input data, blue outlines indicate pre-processing outputs and purple outlines indicate standard-space (MNI152 2mm) data. Rectangular blocks indicate functions and parallelograms indicate transformation objects. In *T*_1_ space, we filtered the *T*_1_ map with a Gaussian median filter to correct for noise caused by numerical instabilities in the *T*_1_ reconstruction from the two MP2RAGE inversion images. We used the second inversion image to mask the brain and obtained probabilistic tissue masks from the brain-extracted *T*_1_ map. We registered the MNI152 *T*_1_ template to the brain-extracted *T*_1_ map and applied the resulting transformation to the MNI152 ventricle and cerebellum mask. In EPI space, we corrected the SE-EPI BOLD scan for motion based on the brain region. We combined the full motion-corrected image, including non-brain tissue, with the top-up scan to correct for distortions along the phase-encoding direction. Based on the corrected image, we extracted the brain. We registered the *T*_1_ map to the SE-EPI BOLD scan and applied the resulting transformation to all *T*_1_ space masks. The brain mask was then thresholded (at 0.9) to reduce partial voluming. We multiplied this brain mask with all images. We binarized the WM, GM, and CSF segmentation by assigning the most probable tissue type to each voxel. Lastly, We included all nonzero probability voxels in the ventricle and cerebellum mask.

#### D.4 SE-EPI data inspection

**Figure D.5.**
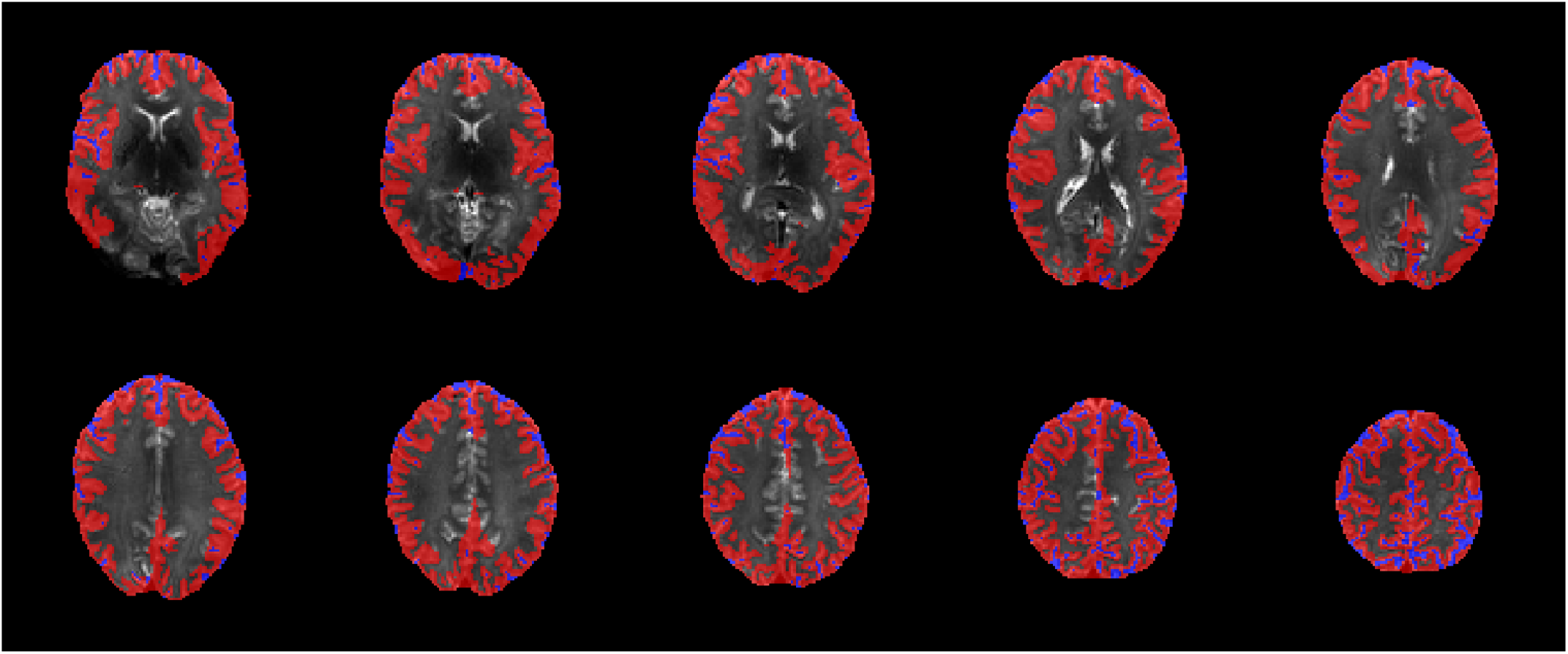
GM (red) and CSF (blue) ROIs overlaid on the corresponding SE-EPI intensity image (ETL=63). Transversal slices are shown at equidistant spacing across the FOV in the feet-head direction. The ventricles and cerebellum were successfully removed from these ROIs, leaving cortical gray matter and sulci-located CSF.

**Figure D.6.**
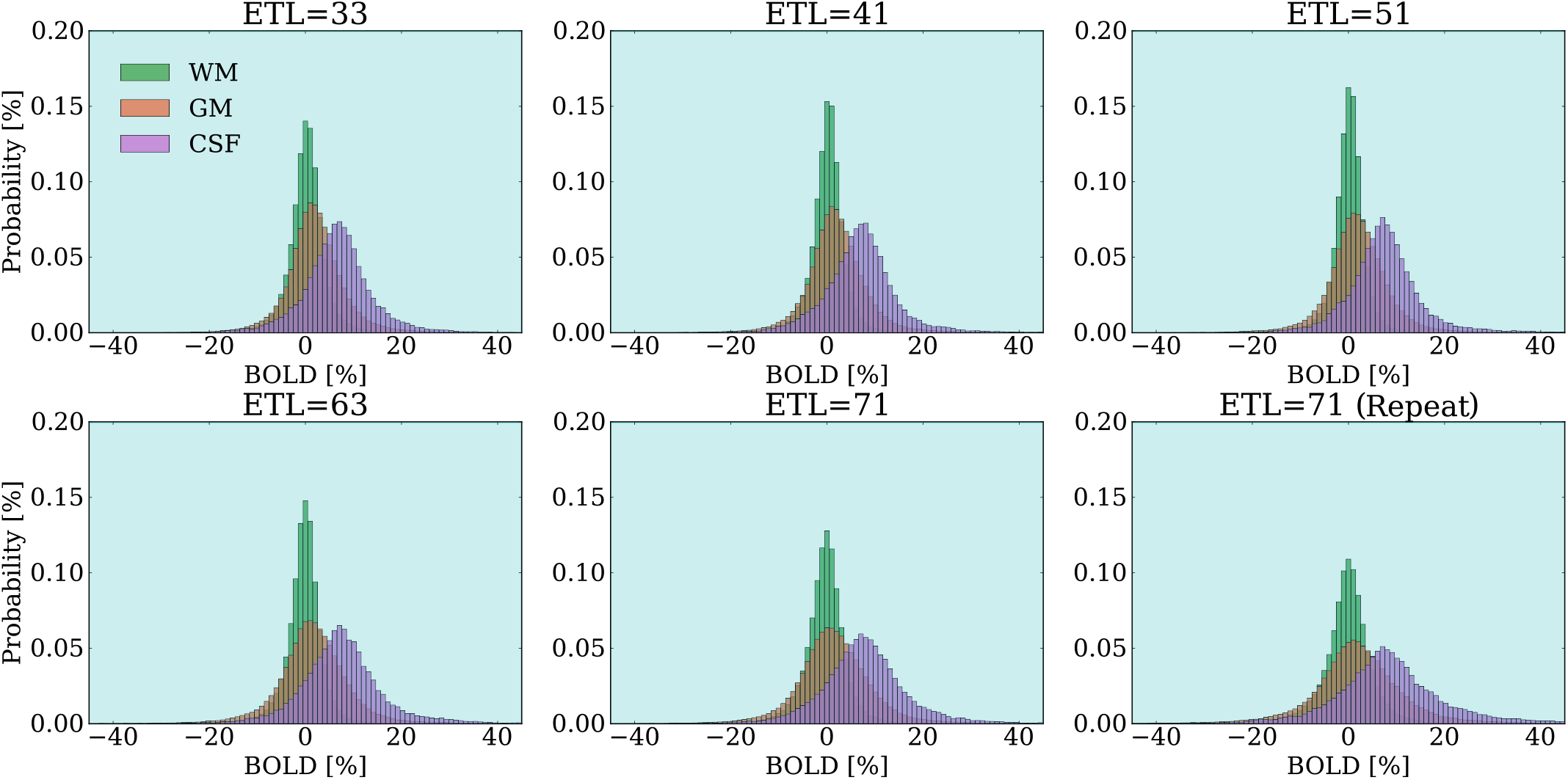
BOLD value histograms for all the acquisitions. We selected only the activated state dynamics (indicated with cyan). We fixed the bins to a width of 1 and a range of -45 to 45. Outside of this range, outliers are present. It is visible that BOLD values within the GM and CSF ROIs are approximately Gaussian distributed. We, therefore, expect that using the median does not bias our estimations of the ‘average’ voxel behavior. The GM ROIs consisted of 49,915 to 50,456 voxels, whereas the CSF ROIs consisted of 6,227 to 6,303 voxels.

**Figure D.7.**
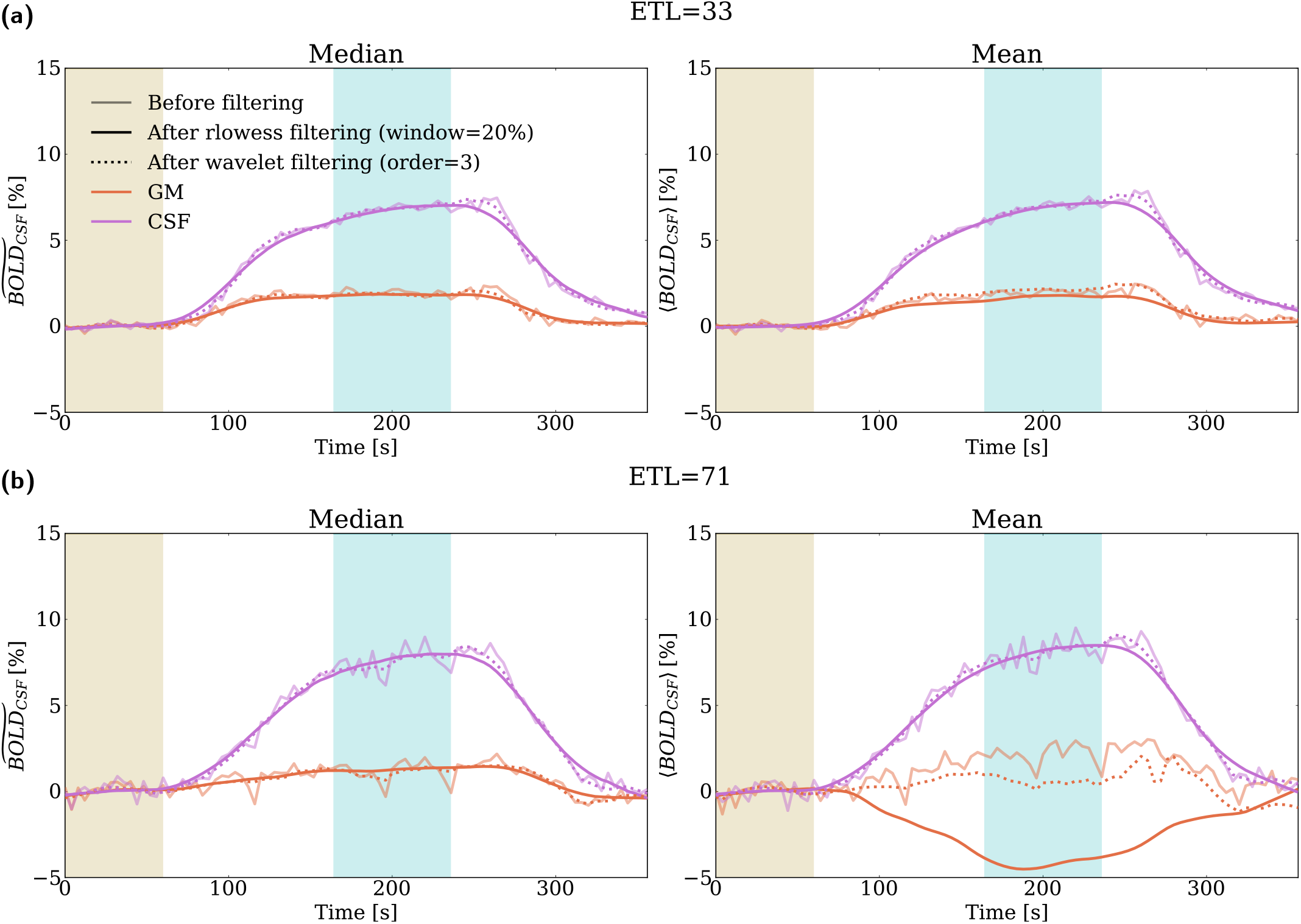
%BOLD responses upon using the ensemble median (indicated by superscript *∽*) versus mean (indicated by brackets <>) as a measure of central tendency. Yellow shading represents the baseline period, and cyan shading represents the activated period. The mean and median give similar results for acquisitions with naturally few outliers and fluctuations, such as ETL=33, shown in panel (a). In contrast, the mean shows high sensitivity to the smoothing method and settings for noisier acquisitions, such as ETL=71, shown in panel (b). These strong effects hint at a dominance of a few outliers. The median is much more robust in these cases. Furthermore, we compared our temporal filtering method rlowess to an alternative wavelet denoising method. rlowess is a robust filtering method and yields smoother BOLD responses in the activation period than the wavelet approach for these settings.

**Figure D.8.**
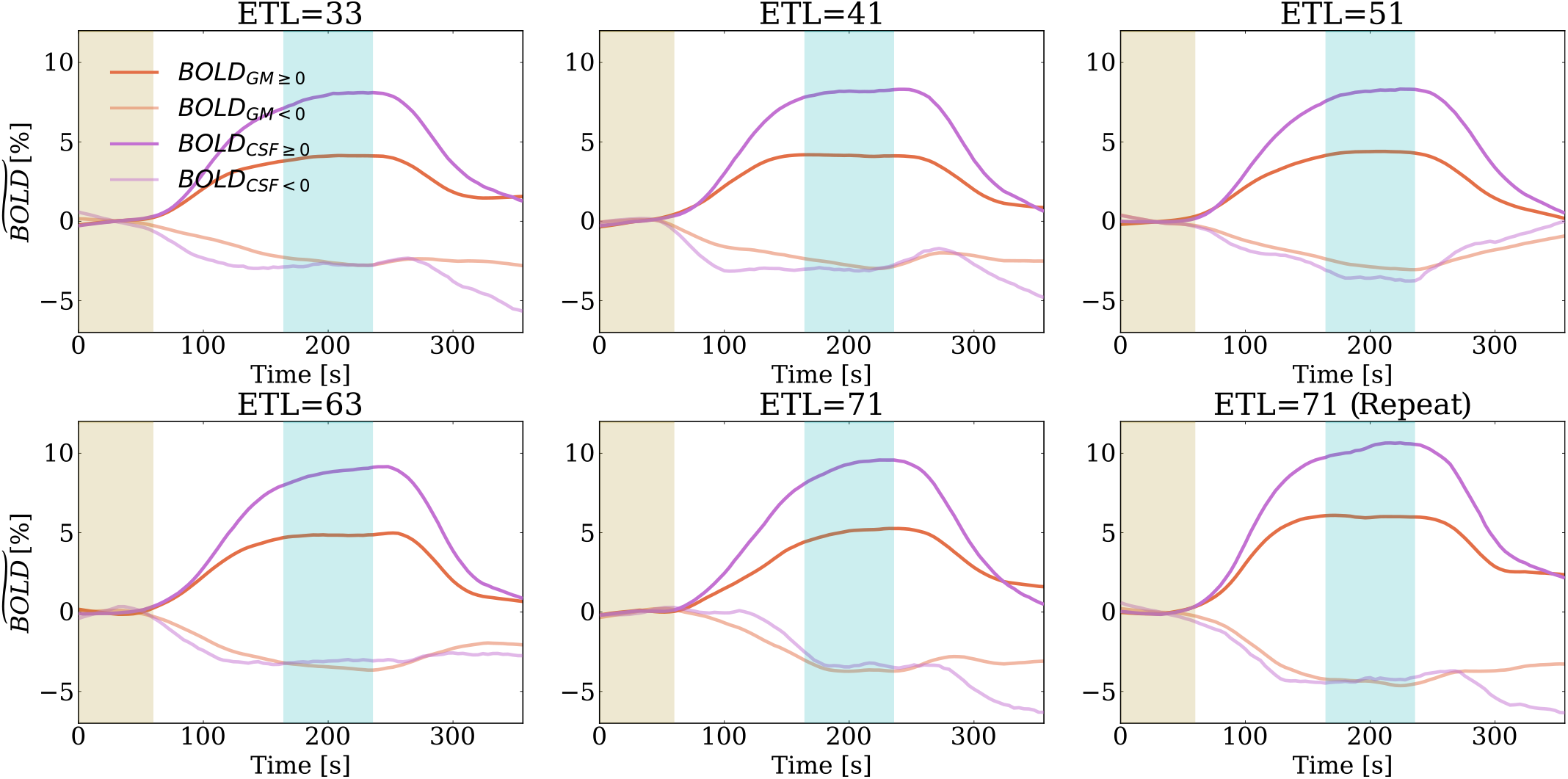
GM and CSF %BOLD responses in voxels with an on average positive (*≥*) and negative (<) BOLD contrast. Yellow shading represents the baseline period, and cyan shading represents the activated period. Negative BOLD contrast was prevalent in our scans: in GM, about 40% of all voxels showed a negative BOLD response, whereas, in CSF, this was about 20% of all voxels. The median negative %BOLD curves showed a characteristic response to the hyperoxic stimulus, which hints at a non-random physiological effect driving the negative BOLD contrast.

**Figure D.9.**
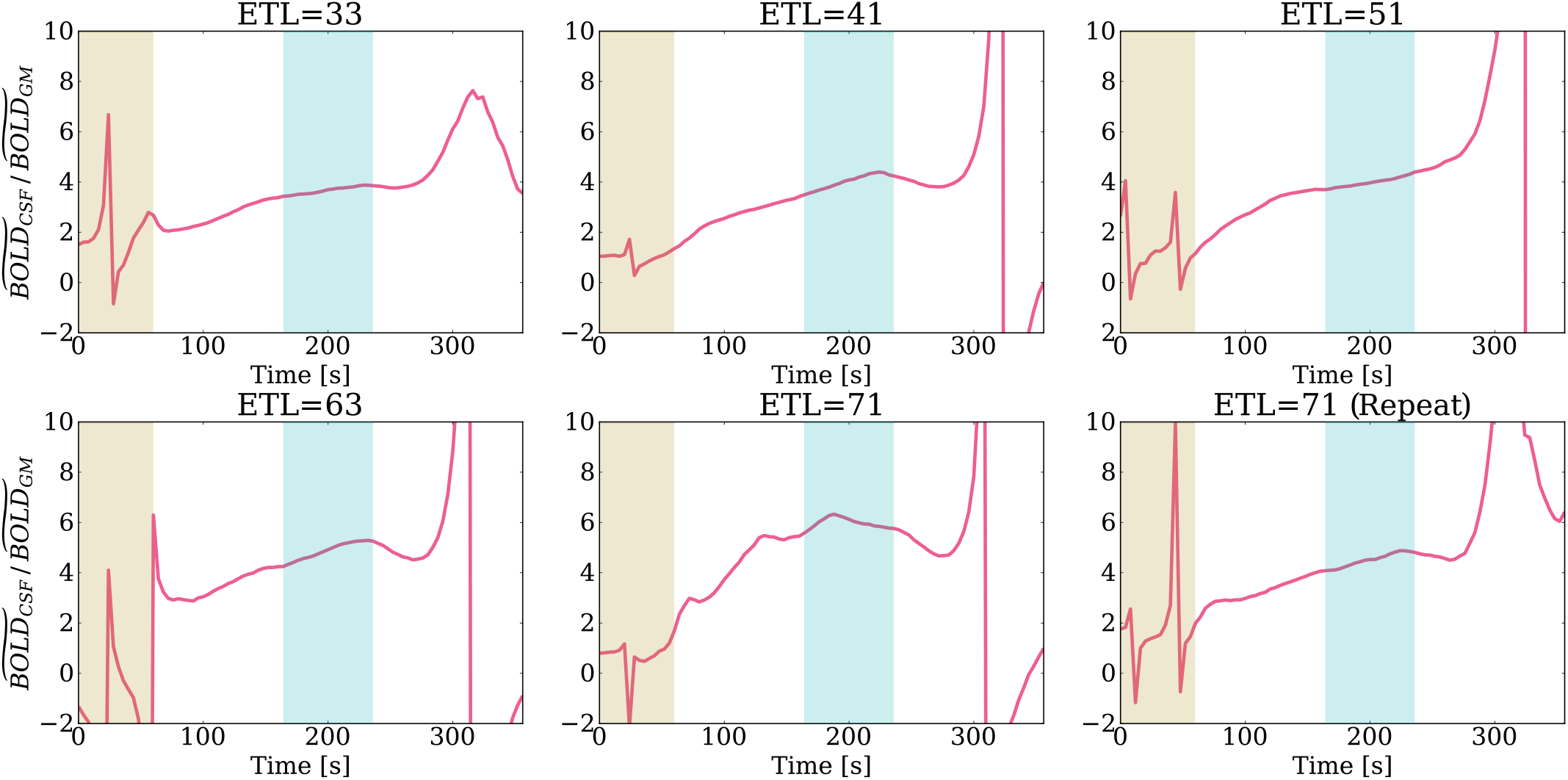
Time courses of the CSF/GM %BOLD ratio for all the acquisitions. Yellow shading represents the baseline period, and cyan shading represents the activated period. In the baseline and recovery period, the ratios fluctuated heavily and diverged as the %BOLD values approached zero. During the activated period, the ratios were relatively stable, although most acquisitions showed a slight increase in CSF/GM %BOLD ratio during the activated period.

**Figure D.10.**
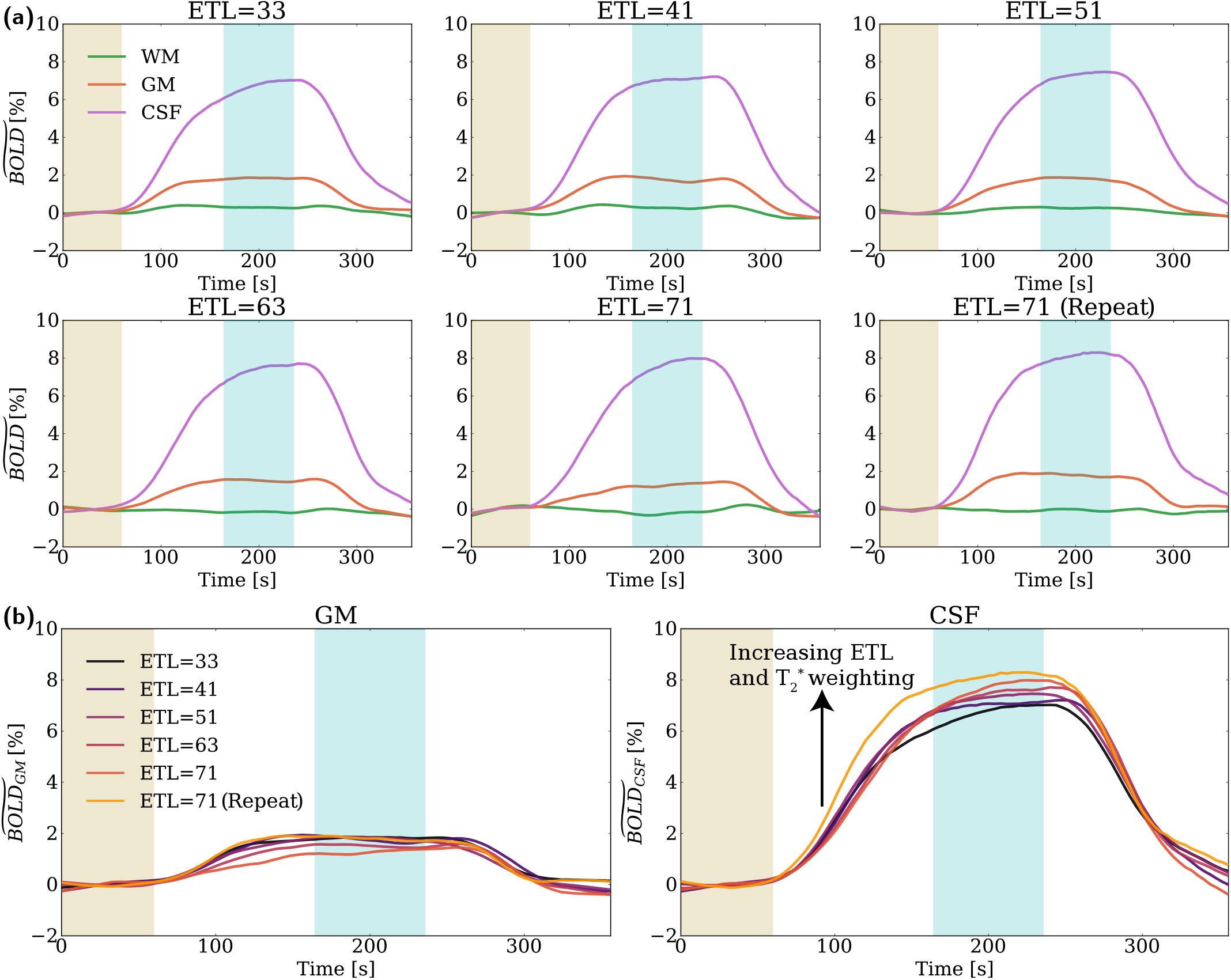
Measured median %BOLD responses for each ETL acquisition in the GM, CSF, and WM ROIs. Yellow shading represents the baseline period, and cyan shading represents the activated period. Panel (a) shows the ROI time traces per acquisition. All acquisitions showed a distinct BOLD response in GM and CSF during the hyperoxic stimulus. The measured GM and CSF %BOLD responses in the activated state were in the same order of magnitude as our simulated %BOLD signal changes (Figure 5, panel a). The measured WM %BOLD responses were close to zero, which we expected considering the poor vascularization and low blood flow in this brain region. Panel (b) shows overlays of the GM and CSF %BOLD responses. The GM %BOLD response tended to decrease as a function of ETL, whereas the CSF %BOLD response increased as a function of ETL. The consistent increase in the CSF %BOLD response as a function of ETL agrees with the simulated trend in Figure 5(a). Note that the ETL=71 repeat scan had an uncharacteristically high GM %BOLD response and a significantly different CSF %BOLD response than the other scans. For this reason, we focused on the first ETL=71 acquisition when we assessed the CSF to GM %BOLD ratio as a function of ETL.

#### D.5 RespirAct data inspection

**Figure D.11.**
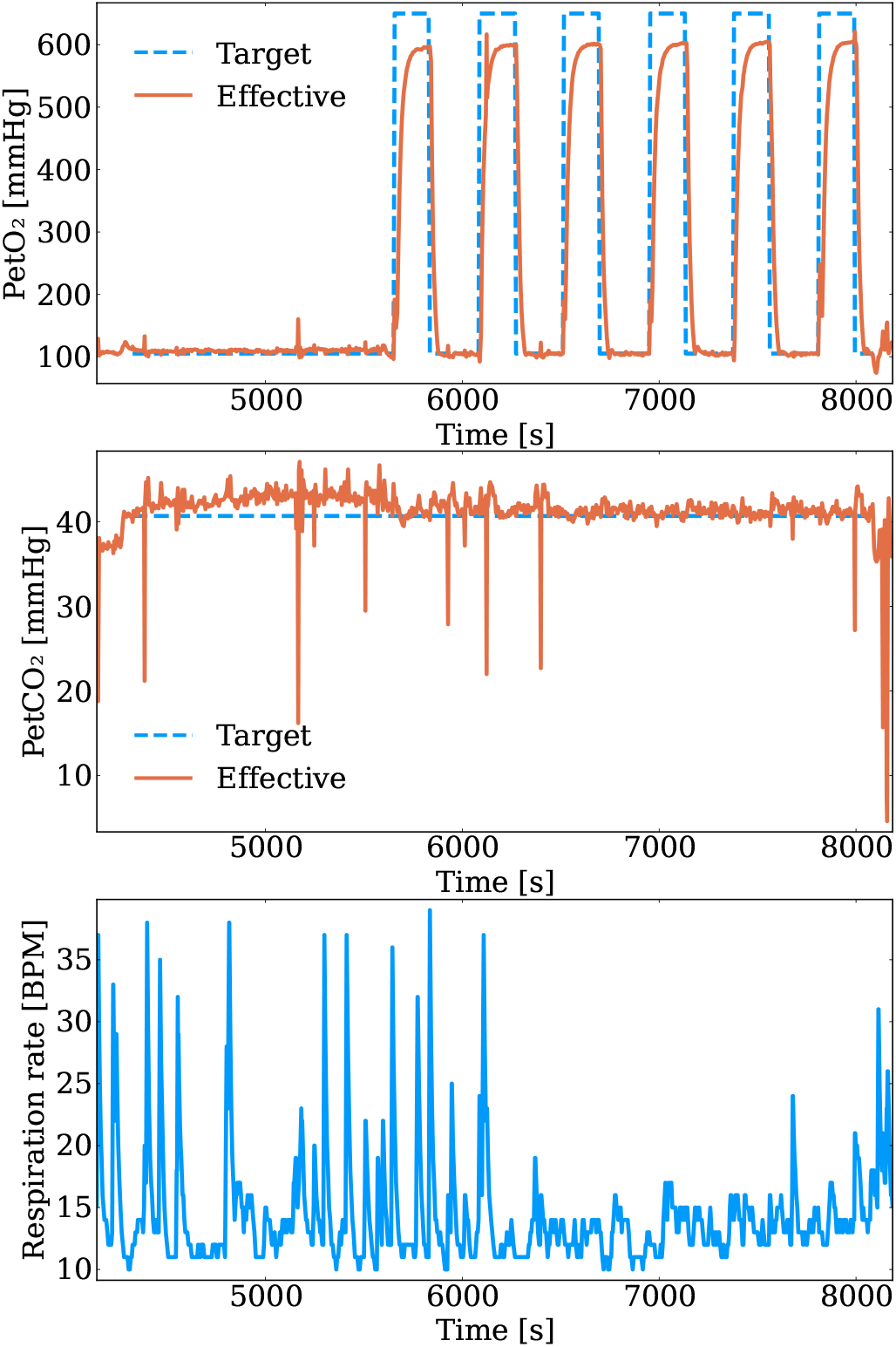
Physiological data from the RespirAct system. The end-tidal CO_2_ pressure (PetCO_2_) was targeted at a constant level of 40 mmHg during all phases of the hyperoxic stimulus. The effective PetO_2_ and PetCO_2_ traces show a peak respectively dip in an early phase of the second stimulus (ETL=71 repeat scan). The respiration rate, given in breaths per minute (BPM), is relatively unstable during the first two (ETL=71) acquisitions and stabilizes afterward.

